# Deciphering a common code: Unitary gene profile factor enabling Spontaneous Tumor Regression

**DOI:** 10.64898/2025.12.20.695084

**Authors:** Arushi Misra, Bindu Kumari, Prasun K. Roy

**Author notes:** Corresponding author’s address: Prasun Kumar Roy, R-206, Dept. of Life Sciences, Shiv Nadar University, Delhi NCR, Dadri 201314, India.

## Abstract

Spontaneous-remission of Cancer (SRC), well documented in humans, is the paradoxical process of natural permanent tumor elimination by host-tissue system without any toxicity, recurrence or cancer stem-cell formation. This rare biological phenomenon is usually associated with immune activation, apoptosis, tumor microenvironment as well as oncogenic suppression. Thus, to understand this strikingly complex event, it is utmost important to look into the genetic and molecular landscape in-depth. Majorly, cancer mortalities occur due to epithelial tumors, hence, to find out the contributing pathways involved we interrogated epithelial malignancies: Neuroblastoma (neural-crest tissue), and Melanoma (skin tissue). Neuroblastoma, the most prevalent tumor type in newborns, arises from embryonic nerve cells and have a very diverse clinical trajectory. Similarly, Melanoma another common skin tumor, developed from malignant transformation of melanocytes (cells producing melanin pigment). It is well-known that in numerous patients, both these malignant tumors spontaneously regress permanently, while in others, these tumors escalate to fatality. We investigated microarray expression of spontaneous regression of malignant neuroblastoma tumors (n=498), the patients having 4 temporal stages (S1-S2-S3-S4), and malignant melanoma tumors (n=22) having 4 time points (T1-T2-T3-T4) [i.e. total 520 patients, each 8 remission-route steps]. We studied differentially expressed genes (DEGs) common in both regressing Neuroblastoma and Melanoma, and analyzed the protein-protein interactions (PPI) as well as protein-miRNA networks. One crucial ubiquitous gene factor was found across each stage of both the regressing tumors. This focal gene codes for a spectrin-repeat protein which acts as a linker between nucleus-cytoskeleton, and aids in maintaining normal cell’s nuclear and cellular integrity (demarcated as a significant morphological change in tumor cells) by abetting the formation of Mut-S complex, which is a significant player in mis-match repair pathway. Strikingly, both this protein and miRNA were found to be commonly targeting PIK3CA protein through separate paths. We endeavor to unravel the mechanism of action and decipher the common gene factors across the entire tumor regression process in both Neuroblastoma and Melanoma. We meticulously identified the underlying biological pathways responsible for spontaneous regression. These results provide insights into the molecular genetics and underlying pathophysiology of SCR in Neuroblastoma and Melanoma, suggesting a role for precision therapeutics in the future that can help mimic this extraordinary prodigy by targeting the uniform factors mentioned in this study.

## 1. INTRODUCTION

The process of spontaneous remission or regression of cancer is the process of permanent tumor elimination by the body’s tissue system and is well documented clinically in humans. For instance, in population-based cancer cohorts tracking, e.g. the Scandinavian **[1]** and Wisconsin **[2]** cancer registries, it is found that 28-32% of malignant breast tumors undergo permanent spontaneous remission endogenously, without any treatment intervention. Spontaneous-remission of Cancer (SRC), well documented in humans, is the paradoxical process of natural permanent tumor elimination by host-tissue system without any toxicity, recurrence or cancer stem-cell formation. Investigation into SRC phenomenon and associated molecular pathways/targets has much innovative prospects, possibly discovering novel biomolecules that may duplicate SRC process on cancer patients. Moreover, some considerable disadvantages of cancer treatment, namely drug side-effects, leftover minute cancer cell population post treatment and cancer stem cells are completely absent in spontaneous tumor regression process.

Neuroblastoma (NB) is a pediatric tumor arising from sympathetic nervous system **[3]**. It affects almost 1 in 7000 live births, making it the most prevalent extracranial solid tumor in children **[3–5]**. The Children’s Oncology Group (COG) has classified NB patients into three risk categories: low, intermediate, and high **[5]**. Factors like the age at diagnosis, MYCN amplification status, International Neuroblastoma Staging System (INSS) stage, histology, and tumor cell ploidy are used to classify this risk **[5,6]**. Melanoma spreads locally, regionally, and distantly, which sets it apart from nonmelanoma skin malignancies. The degree of invasion and ulceration of the initial lesion directly affects a person’s risk of metastasis. Invasion, angiogenesis, extravasation, dispersion, and colonization of the target organ are all part of the early phases of cancer metastasis. Primary cutaneous melanoma is still the most deadly type of cutaneous tumor, and its frequency has been rising continuously for several decades. Melanoma has a good survival rate of about 94% if detected in its early stages **[7]**. The National Cancer Institute (NCI) estimates that the incidence of metastatic melanoma was 0.9 per 100,000 between 2014 and 2018. The prognosis is usually worse for ocular and mucosal melanomas **[8]**. Melanoma regression, often referred to as fully regressive melanoma, is a condition in which the initial cutaneous melanoma is completely replaced by fibrotic components as a result of the host immune response.

Differential gene expression of thousands of genes in biological samples can be monitored simultaneously through the use of DNA microarrays for gene expression analysis **[9,10]**. In addition to identifying genes that may be of prognostic value **[11]**, as well as genes that might be linked to aggressive tumor behavior **[12]**, microarray gene expression analysis offers new insights into the genetic basis of tumor initiation and progression. Furthermore, the analysis of tumor sensitivity to chemotherapy or other targeted therapeutic methods has demonstrated potential with microarray gene expression profiling **[13, 14]**. Maneuvering differential gene expression analysis of the microarray datasets corresponding to Neuroblastoma and Melanoma available at Gene Expression Omnibus, we could overview an entire genetic landscape of the SCR phenomenon occurring in both the tumors discussed in this study. Thereby, we could formulate a uniform gene expression pattern and the mechanism underlying spontaneous cancer regression. This study discusses the crucial importance of the common gene factor SYNE1 along with the protein it codes for that is Nesprin-1, in the mechanism of SCR.

Nesprin-1, the largest protein of the nuclear envelope (NE), interacts with Lamins, SUN, Emerin and Actin proteins to form a network along the NE. Numerous age-related illnesses, as well as cancer, have been linked strongly to mutations in the SYNE1 gene and decreased amounts of the Nesprin-1 protein **[15]**. It has been reported that mutation in the SYNE1 gene that results in decreased Nesprin-1 protein levels, triggering altered nuclear shape, altered amounts and localization of NE components, centrosome localization, and genome stability, which indicates that Nesprin-1 overexpression could reverse the malignant phenotype of these cells.

One may note that, these SCR processes are actually insightful, since normal tissue is protected during the tumor regression, and cancer stem cells are also eliminated, there being full recovery of the patient. Here, we investigate such SCR process in tumors of different tissues and thereby endeavor to unravel the commonalities of the factors that may be resulting in the SCR process. Thereby we arrive at candidate biomolecules that have the potentiality to induce SCR in different tumors ubiquitously. Furthermore, investigation into this phenomenon and the associated molecular pathways/targets may have much prospects, and these may lead to discovery of novel biomolecules that may duplicate the SRC process on cancer patients, as an innovative therapeutic approach.

## 2. METHODS AND MATERIALS

### 2.1 Data acquirement

We have analyzed the raw microarray preparation of the spontaneous tumor regression model of melanoma bearing Libechov minipigs (MeLiM) from the ArrayExpress platform (https://www.ebi.ac.uk/biostudies/arrayexpress/studies/E-MEXP-11522query=E-MEXP-1152) in our earlier study **[16]** and in this current study, we have analyzed the processed data of the Agilent microarray datasets of humans GSE49710 (n = 498) from Gene Expression Omnibus (GEO) database-NCBI (National Center for Biotechnology Information). The genes expression levels were already processed and log2 transformed and further downloaded using R-studio platform. In case of Melanoma, the tumor spreads and kills about half of the subjects while in the case of Neuroblastoma, about 21% of the patients died and rest stayed alive. Nevertheless, in the other half of animals in Melanoma as well remaining 79% of the patients of Neuroblastoma, the tumor develops up to a certain point (tumor progression) and then undergoes spontaneous shrinking and healing (tumor regression), and these latter animals and patients remain healthy thereafter.

Normalisation and Differential expression analyses were performed by “limma” package using the R (version 3.6.2) software for Melanoma and by “limma” and subsequently “EdgeR” using R (version 4.4.0) for Neuroblastoma respectively.

### 2.2 Identification of differentially expressed genes (DEGs) of spontaneous melanoma and neuroblastoma regression data

After applying the two primary cutoff criteria, (−2=> FC <=+2, p-value <=0.05) and (−0.5 =>FC >= +0.5, p-value <= 0.05) we have obtained differentially expressed genes (DEGs) at the above-mentioned time points t1, t2, t3, and t4 in Melanoma and Stages S1, S2, S3, S4 in Neuroblastoma respectively. For this current study, we have taken three types of approaches:

a. Type-1: For this gene set, we choose the genes whose expression wa s consistent from time points t2 to t4 across the Melanoma regression process, and stages S3 & S4 for Neuroblastoma regression process.
b. Type-2: For this gene set, we now consider those genes whose expression levels differ between time points t1 (full progression) and t4 (full regression), and between stages S1 (full progression) and S4 (full regression); for example, a gene that is upregulated at t1/S1 should be downregulated at t4/S4, and vice-versa.
c. Type-3: This is the major protagonist gene set/gene common across all time-points and stages of Melanoma and Neuroblastoma respectively.

We have also generated the protein-protein interaction (PPI) networks for selected DEGs using String platform. It generates molecular functions of the protein complexes arranged in networks that are consolidated by electrostatic or biochemical factors **[17]**.

### 2.3 Gene Ontology Analysis

To carry out the gene enrichment analysis of DEGs, ClueG0-v2.5.5 and CluePedia platform from Cytoscape **[18]**, the FunRich facility (version: 3.1.3) **[19]**, and the DAVID platform **[20]** (https://david.ncifcrf.gov/) have been utilized. The Gene Ontology (GO) results for cellular component (CC), biological process (BP), and molecular function (MF), and Immunological processes (IP) were ranked by p-value. Therefore, we used the conventional adjusted kappa statistic of > 0.4 to establish and compare the network of functionally linked GO terms in this investigation.

### 2.4 Identification of most significant regression gene(s)

Now, we need to identify the most effective melanoma and neuroblastoma-regressing driver gene profile from the afore-mentioned three types of approaches for selecting DEGs.

Respectively, we have-

i. Firstly, shortlisted common genes between aforementioned Type I set of genes of Melanoma and Type I set of genes of Neuroblastoma.
ii. Secondly, we have selected the common gene among: the spontaneous melanoma and neuroblastoma microarray data genes [i.e. those genes whose expression levels have opposite signs (negative or positive) at time points t1 & t4 and stages S1 & S4 respectively],
iii. Thirdly, we found our only common gene in each time-point of Melanoma and each stage of Neuroblastoma.

### 2.5 Identification of Significant proteins

For majority cancer types, targeted molecular treatments are acknowledged as a revolutionary therapeutic approach. Thus, we investigated the protein that was encoded by the gene that was discovered to be prevalent at each melanoma time point as well as each neuroblastoma stage, which turned out to be Nesprin-1. Thereafter, we investigated the related interacting proteins along with the signaling pathways these proteins are involved in. Recent data indicates that blocking a single effector protein of the signal transduction cascades involved in tumor pathogenesis is insufficient to prevent or stop tumor growth; instead, the most effective targeted tumor treatment requires a multimodal molecular modification and inhibition **[21]**.

Thus, we further delved into the other molecular aspects.

### 2.6 Identification of Significant miRNA

miRNAs are the non-coding RNAs which have been shown to play major roles in all pathways, by either inhibiting or enhancing protein expression by binding to its 3’ or 5’ UTR. To investigate the regulatory relationships and landscape of interactome of identified protagonist gene and miRNAs, three miRNA targets prediction databases were used. The miRNAs predicted by all the databases were shortlisted as the hub/major targeted miRNAs.

The co-expression network based on the correlation analysis between the hub genes and miRNAs was constructed by Cytoscape software.

### 2.7 Common signaling pathway playing role in Melanoma & Neuroblastoma

Recent developments in molecular oncology have produced new treatment approaches that specifically target-

i. immune regulatory molecules involved in suppressing the anti-tumor immune response, such as T-lymphocyte-associated antigen 4 (CTLA4), programmed cell death1 receptor (PD-1), and its ligand (PD-L1), and
ii. key effectors of the pathways; for instance mutations in significant genes, which are found to play an important role in the pathogenesis of melanoma & neuroblastoma **[22,23]**.

Apart from these well-known pathways, we tried to find the pathways commonly targeted by our protagonist gene as well as its interacting miRNA, and surprisingly we found that not only the common gene factor but also its interacting miRNA targets PI3K/AKT pathway, through their respective routes, thereby commonly inhibiting AKT pathway and subsequently cellular proliferation.

Secondly, the protein coded by our common gene factor is involved in Mis-match repair pathway thereby leading to DNA-damage repair, which is explained further in the results and in detail in the discussion section.

The most significant pathways involved with our protein of interest (Nesprin-1)-

i. PI3Kinase-AKT Dependent pathway
ii. Mis-match repair pathway

## 3. RESULTS

In order to address this rare natural phenomenon of spontaneous regression of cancer, here in this study we have investigated the driving genes and the downstream signaling pathways regulated by these genes, for replicating this extinction phenomenon in melanoma as well as neuroblastoma. Apart from the gene and pathway analysis, we have also addressed the interacting non-coding RNAs (microRNAs) which play significant role in regulating proteins and their functions.

### 3.1 Microarray data analysis and identification of differentially expressed gene

The differentially expressed genes (DEGs) were identified based on fold change value and p-value, as mentioned in the methodology. We have tried to tackle our study to investigate the spontaneous regression driving genes by two types of approaches, as discussed in method section 2.2. From the first approach, we have 176 upregulated and 116 downregulated DEGs whose expression values were consistent from time point t2 to t4 and, 276 upregulated and 175 downregulated DEGs whose expression values were consistent in stages S3 & S4. Likewise, from the second approach, we have 213 DEGs whose expression values have differing signs at time points t1 and t4 and 86 DEGs at stages S1 & S4. We also, constructed the Volcano plot, Mean-difference plot as well as a Density plot to mark a reasonable difference between the upregulated and downregulated genes as well as to analyze the gene expression density between Stage 1 and Stage 4 in Neuroblastoma (Fig. 1). Furthermore, we formulated a String network for these DEGs individually (Fig. S1-S3).

**Fig. 1:**
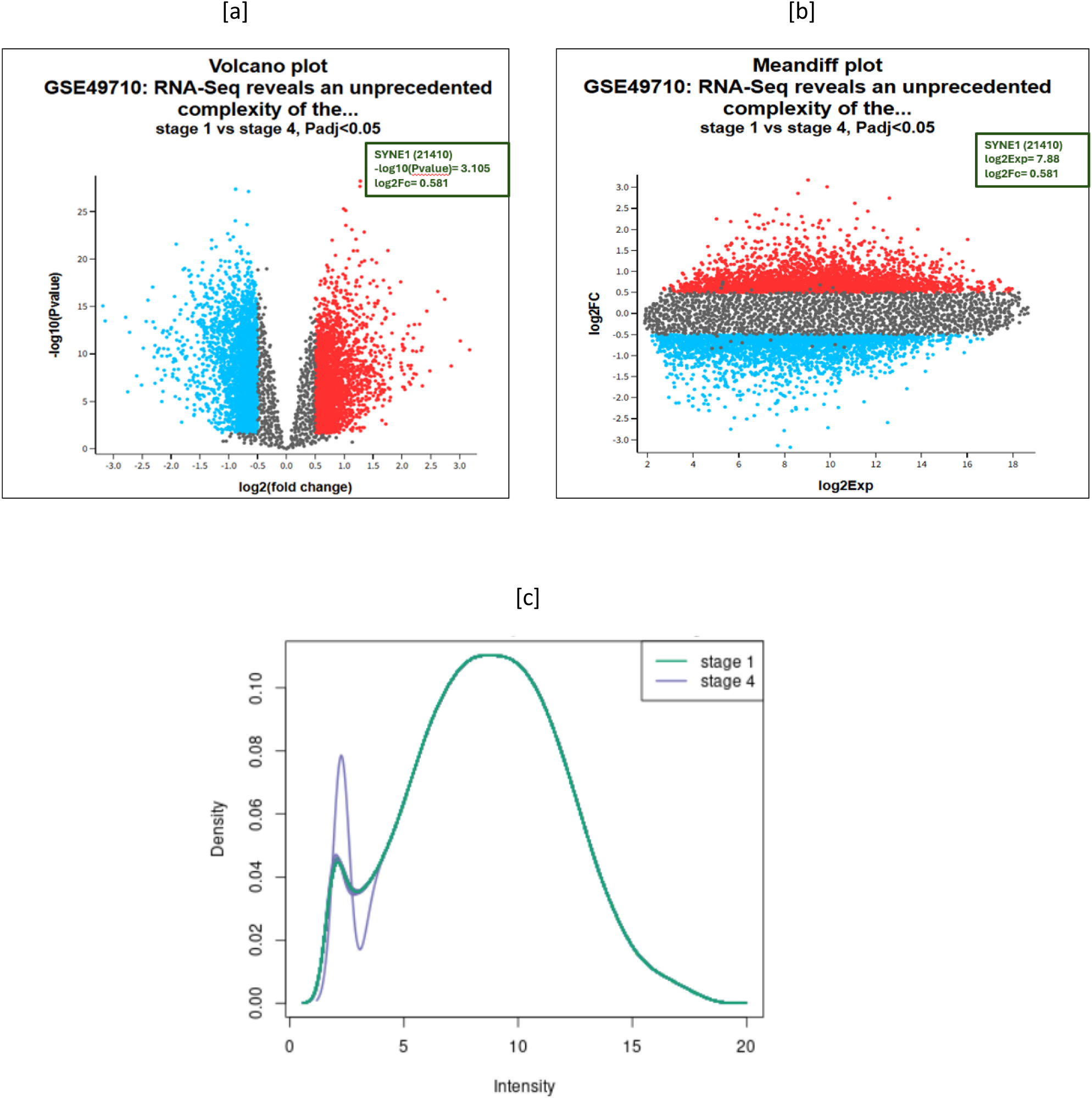
Plots generated between Stage-1 (tumor progressing stage) and Stage-4 (tumor regressing stage) (a)Volcano plot (b) Mean-difference plot (c)Density plot of genes playing role in regression.

### 3.2 Gene enrichment analysis of differentially expressed genes

GO-terms and pathways enrichment were analyzed through multiple databases or software, including PathVisio (https://pathvisio.org/), DAVID (https://davidbioinformatics.nih.gov/), KEGG pathway (http://www.genome.jp/kegg), ClueGO and Cluepedia from Cytoscape (https://cytoscape.org/), FunRich (http://www.funrich.org/) and STRING (https://string-db.org/) platforms with p<0.05 as cut-off criterion. The DEGs narrowed down for Melanoma and Neuroblastoma from two approaches Type I and Type II were subjected to GO analysis and divided into four categories: The Molecular Function (MF), Biological Process (BP), and Cellular Component (CC).

The analysis of the common genes in Type-I and TYPE-II is as follows:

i. In the Molecular function group, the genes were mainly enriched in Cytoskeleton-nuclear membrane anchor activity, Nuclear and cytoskeleton DNA & chromatin binding, actin and protein membrane binding activity (Fig. 2)
ii. Biological processes: the TYPE I upregulated genes were correlated to cell growth and/or development while TYPE I downregulated genes were correlated to cell-cycle, DNA repair, DNA metabolic processes, mitosis, cell communication, and signal transduction. GO term analysis showed that most of the DEGs were enriched in kinetochore, collagen, binding functions, cell cycle, and cell growth (Fig. 3 (a)(b)).The data was in keeping with the knowledge that abnormality of cell cycle and cell growth regulators was the major cause of tumorigenesis. Moreover, the metabolism of the nuclear components and intercellular substances of cancer cells was different from that in normal cells.
iii. Cellular component

**Fig. 2:**
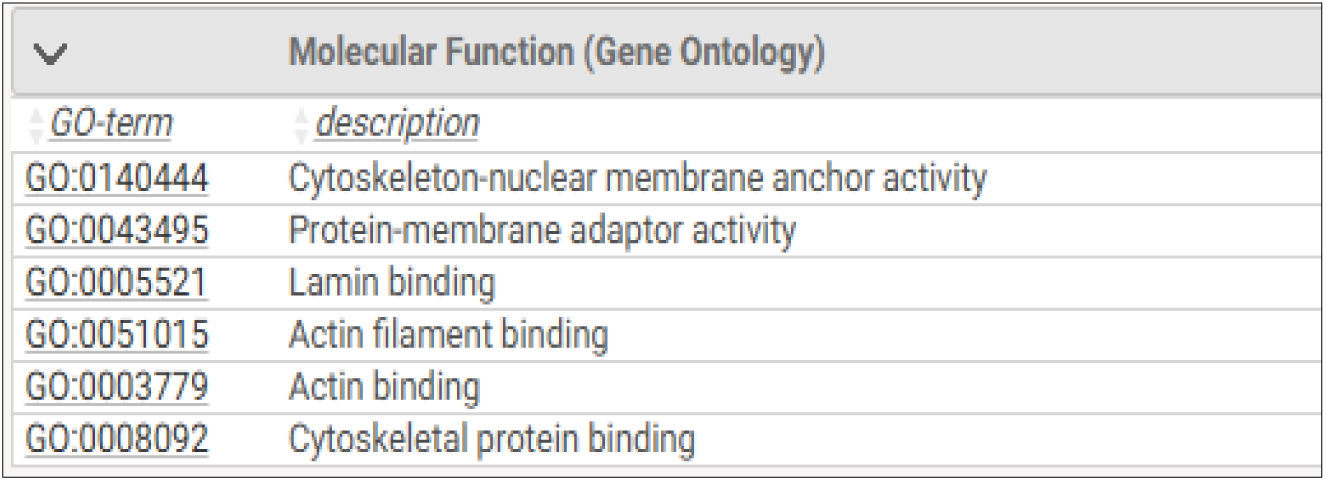
Molecular functions associated with SYNE-1 gene.

**Fig. 3(a):**
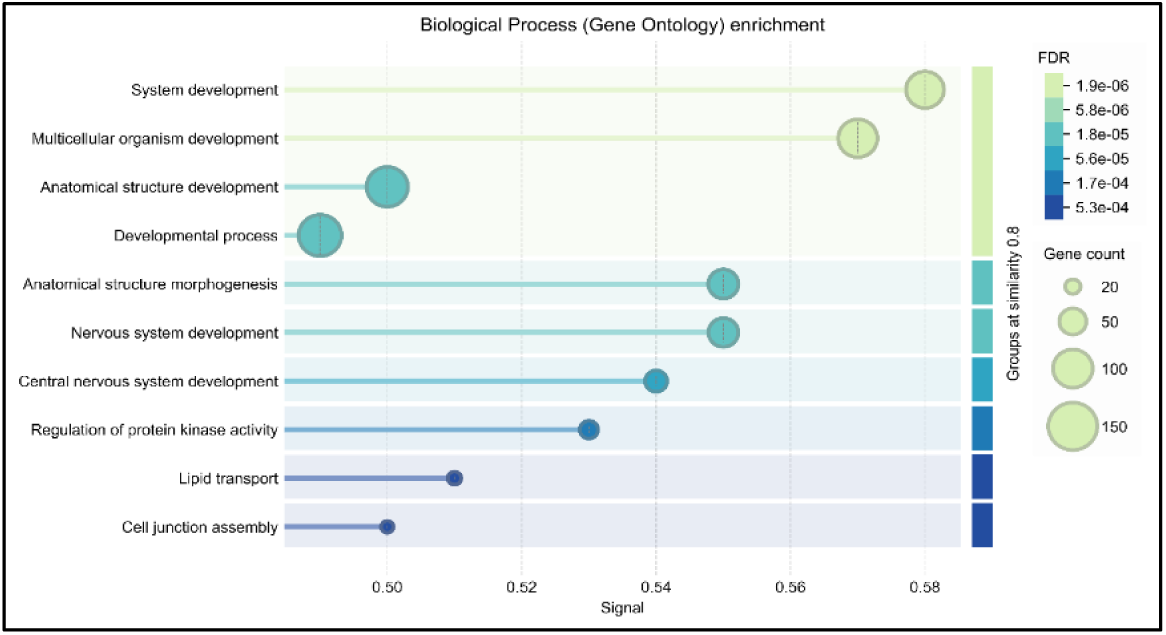
Biological processes enriched in TYPE-I upregulated.

**Fig. 3(b):**
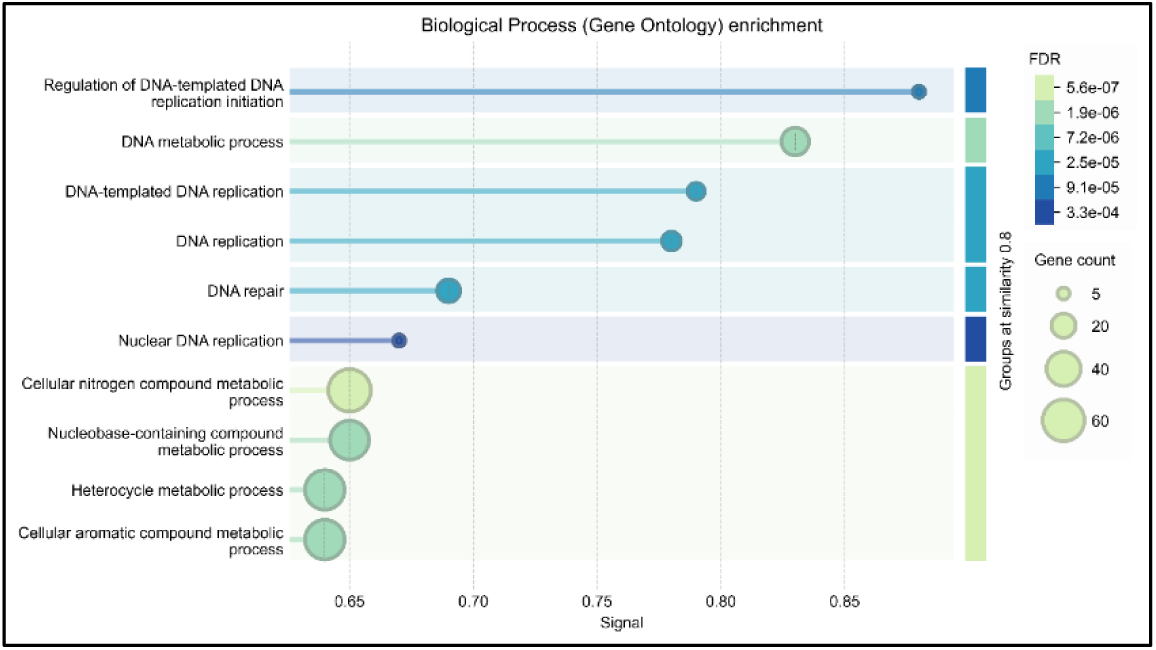
Biological processes enriched in TYPE-I downregulated.

Majorly involves nucleus, nuclear membrane, cytoskeleton filament, lamin filaments, basically indicating nucleo-cytoskeleton conjuncture (Fig. 4)

**Fig. 4:**
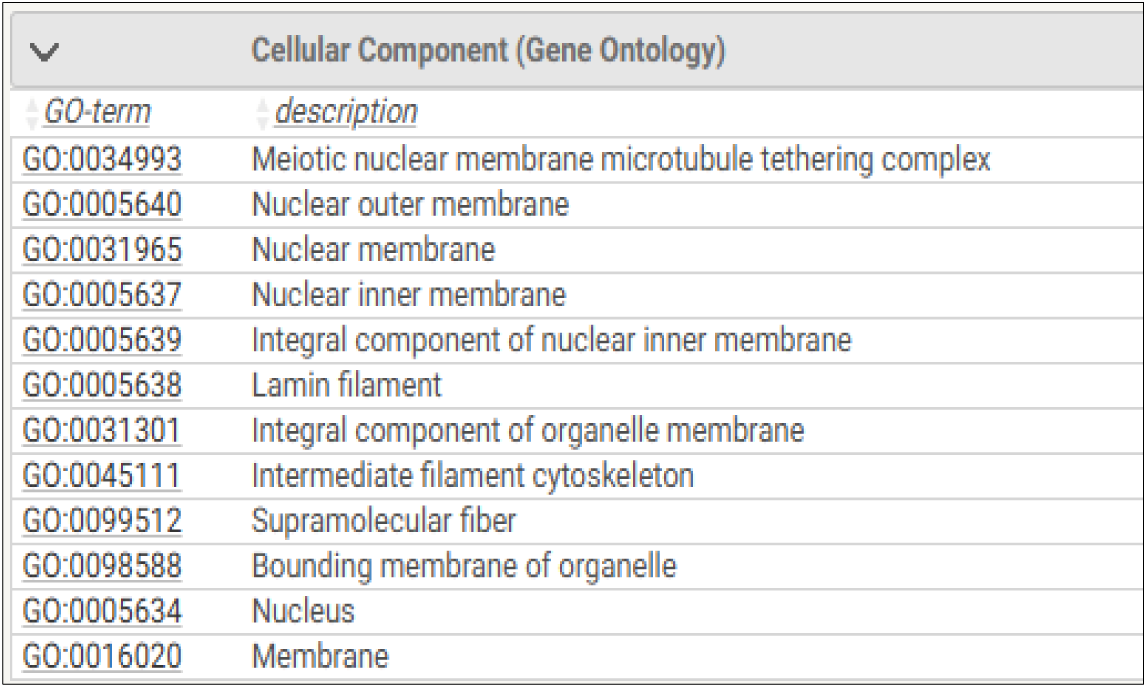
Cellular components associated with SYNE-1 gene.

In order to identify a unitary factor and mechanism for tumor regression in both Melanoma and Neuroblastoma, we looked out for common genes from individual analysis of each of the two tumors.

The genes common in Type I gene set of Melanoma and Neuroblastoma turns out to be a set of 11 genes (Table 1). Out of these, two genes show altering expressions in Melanoma and Neuroblastoma so we omit these and chose the remaining 9 genes as our common hub genes viz-CLU, CRYAB, BCAR3, SYNE1, SMPDL3A, MCM2, CHEK1, C21orf45, FANCD2 which are consistently expressing in the tumor regressing phases of Melanoma as well as Neuroblastoma.

**Table 1:** Type-I hub DEGs common in Melanoma and Neuroblastoma.

| GENES | FOLD CHANGE (logFc) |  |  |  |  |
| --- | --- | --- | --- | --- | --- |
|  | Melanoma |  |  | Neuroblastoma |  |
|  | T2 | T3 | T4 | S3 | S4 |
| CLU | 1.969244026 | 3.64199902 | 2.71229739 | 0.460929802 | 0.987432055 |
| CRYAB | 1.790591444 | 3.47941402 | 3.39478774 | 0.820020094 | 1.055733268 |
| BCAR3 | 0.749077514 | 2.90617844 | 1.20455975 | 0.395686166 | 0.679753466 |
| SYNE1 | 1.109263618 | 2.4731596 | 2.07258008 | 0.622530058 | 0.588639515 |
| SMPDL3A | 1.105790732 | 2.55787255 | 2.46369835 | 0.343908756 | 0.610783026 |
| NRXN3 | -1.021559342 | -1.63262697 | -1.26069186 | 0.83418006 | 0.580611819 |
| MCM2 | -1.12336248 | -1.94399648 | -1.95353705 | -0.535195084 | -1.21271031 |
| CHEK1 | -1.22006972 | -1.5192305 | -1.09156166 | -0.594527559 | -1.24797806 |
| C21orf45 | -1.472043087 | -2.03078431 | -1.20443337 | -0.29237902 | -0.71477061 |
| ECE2 | 1.434572366 | 2.67154152 | 2.0258701 | -0.388529776 | -0.94895403 |
| FANCD2 | -1.332417101 | -1.9884613 | -1.51438986 | -0.52523273 | -1.07282255 |

To further detail in the Biological processes, Molecular functions and Pathways involved of these 9 genes, we analyzed them using ClueGo and CleuPedia from Cytoscape (Fig 5 (a)(b)(c)).

<u>Biological process.</u>

**Fig. 5(a):**
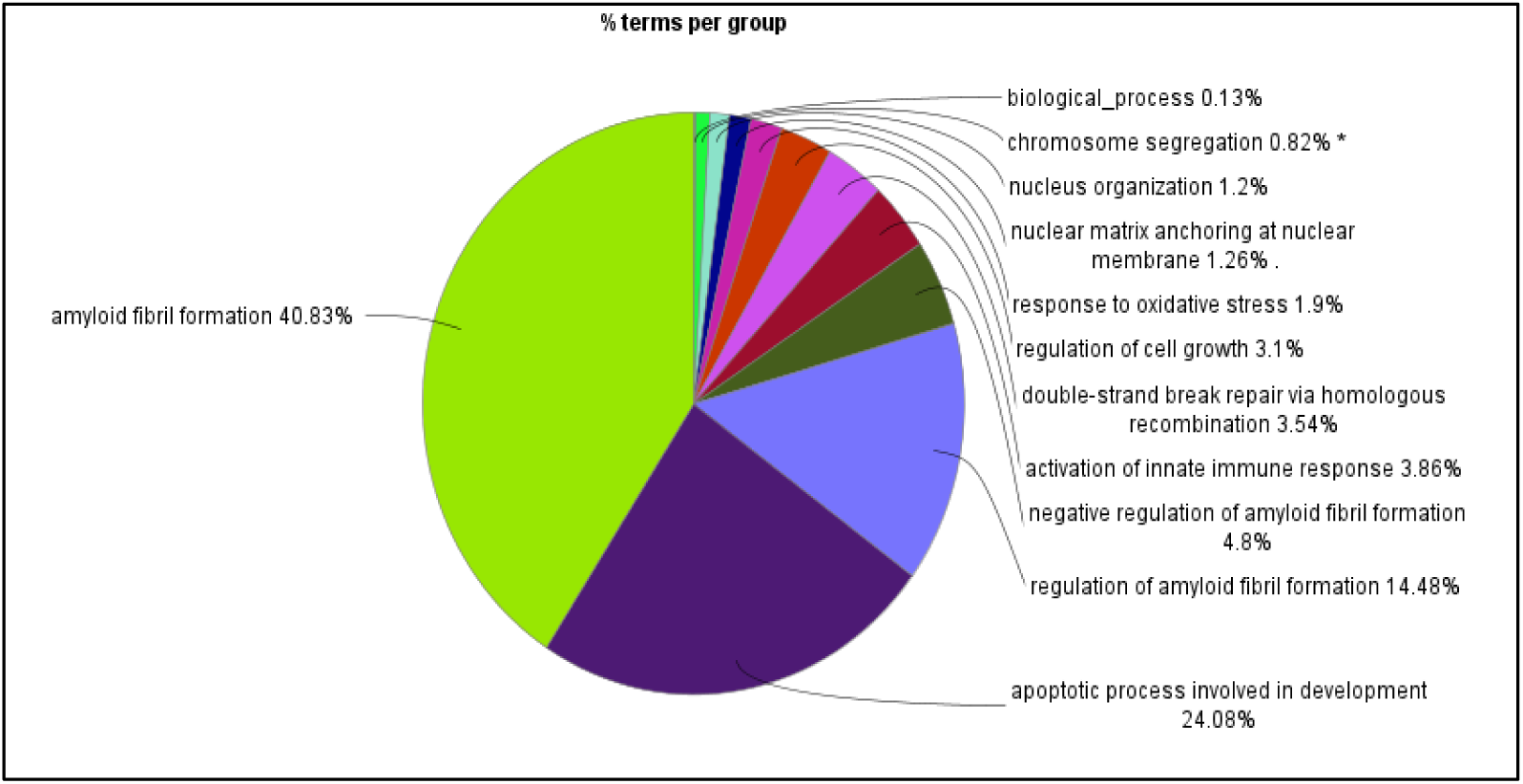
highlights biological processes like nuclear organization and nuclear matrix anchoring.

<u>Molecular functions-</u>

**Fig. 5(b):**
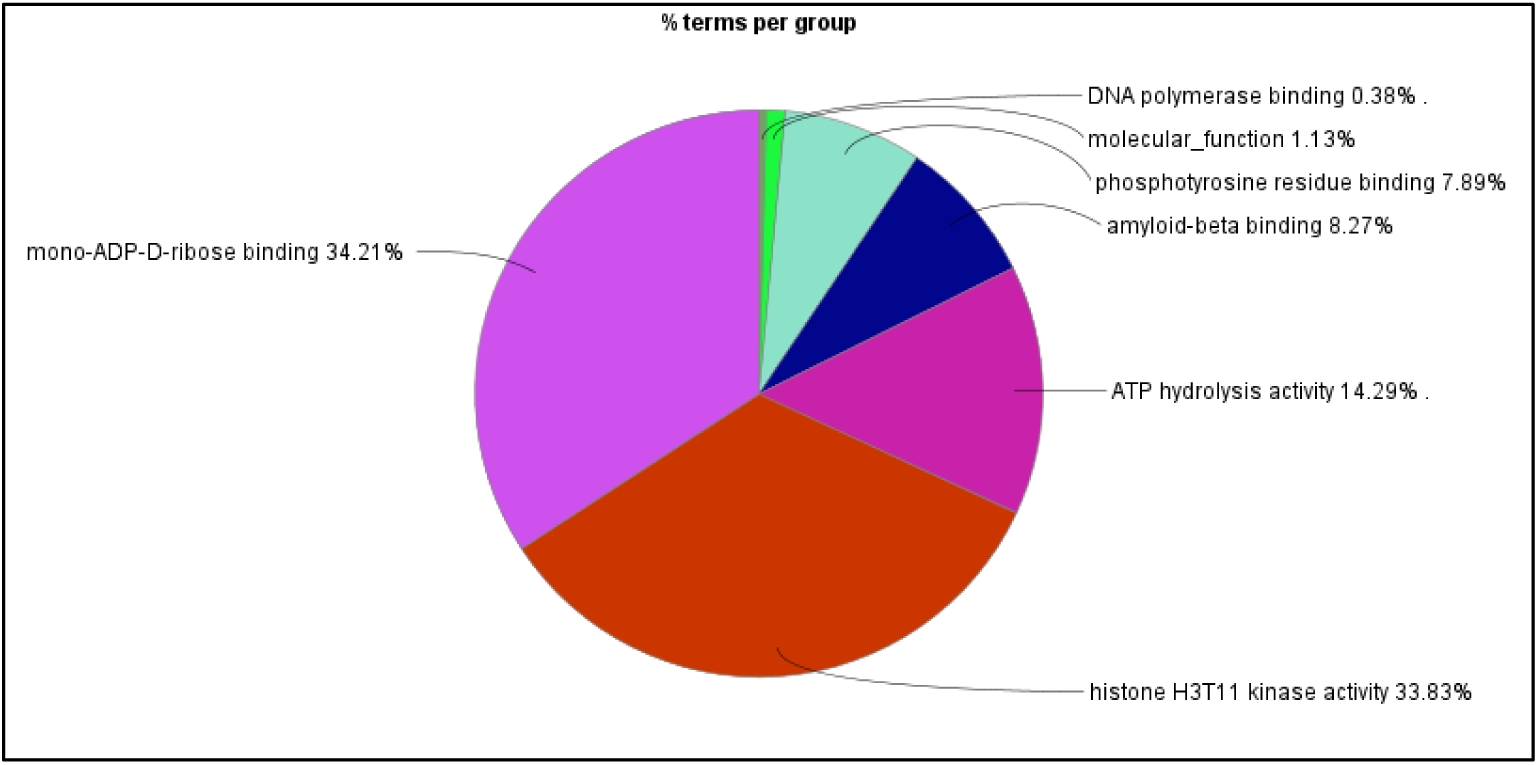
highlights molecular functions like DNA polymerase binding and amyloid beta binding.

<u>Pathways-</u>

**Fig. 5(c):**
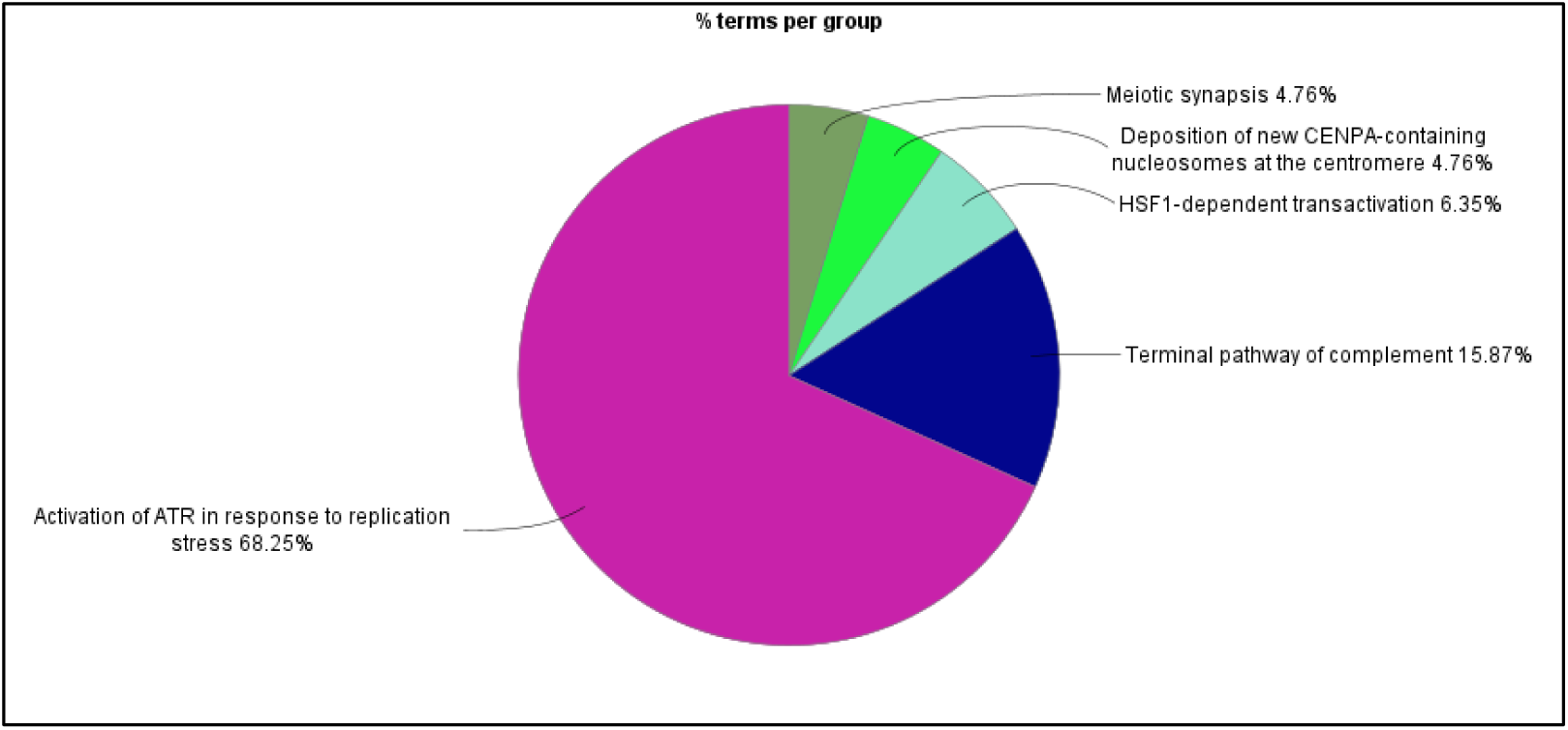
highlights ATR response activation for DNA repair as the major.

Further, by analyzing the TYPE-II set of genes in Neuroblastoma, 29 genes were found to be upregulated in Stage1 and downregulated in Stage 4, while 57 genes were found to be upregulated in Stage 4 and downregulated in Stage 1. In melanoma, a total of 17 genes were found in Type-II analysis. We found a common set of genes in Melanoma and Neuroblastoma from TYPE-II approach and investigated into the Biological processes, Molecular functions, Cellular components and Pathways involved using ClueGO and CluePedia from Cytoscape.

For Neuroblastoma, Stage 1 and Stage 4 were analyzed in opposite combinations (Fig. 6 (a)(b)(c)(d)(e)), as follows-

**Stage 1 downregulated & Stage 4 upregulated genes**-

These genes were found to be aiding the tumor regression process.

<u>Biological processes:</u>

**Fig. 6(a):**
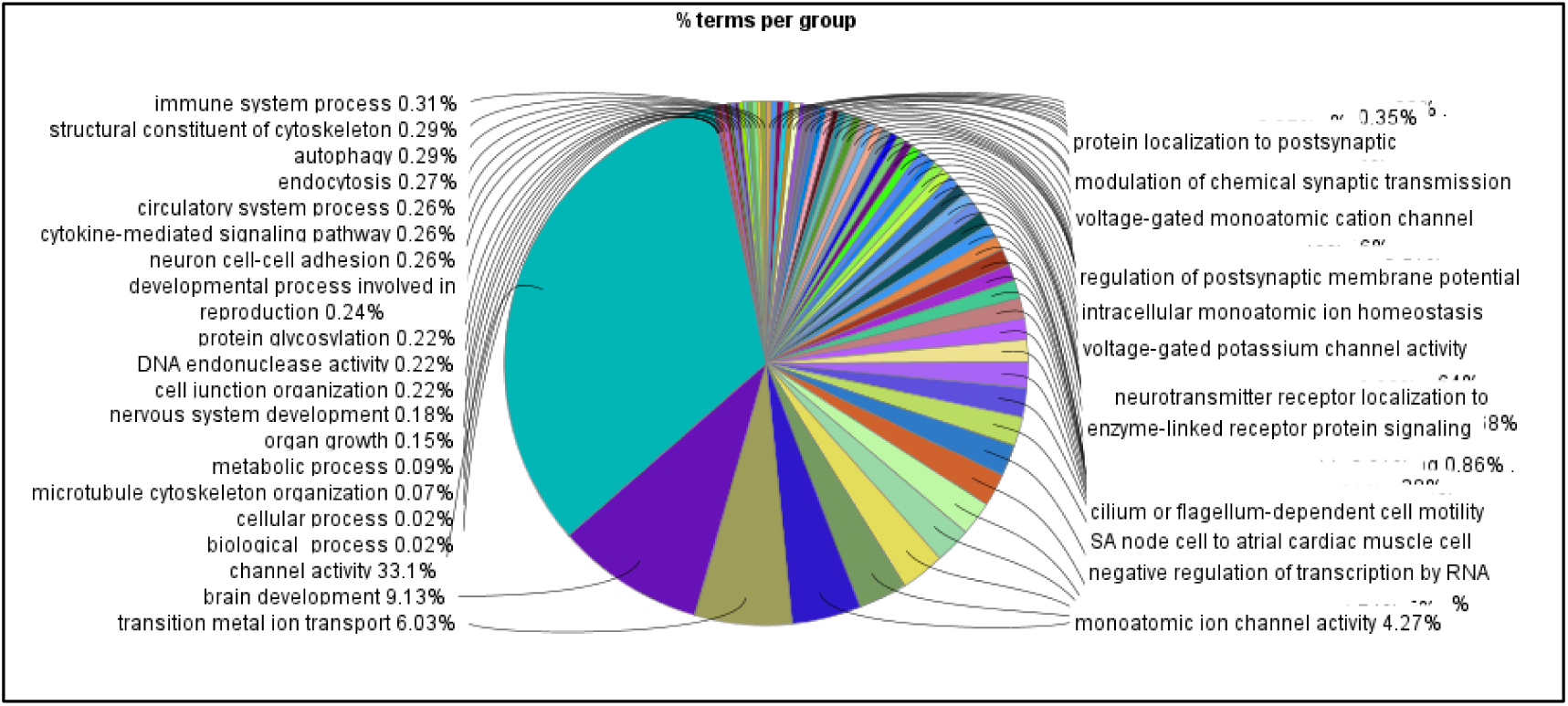
shows pie chart highlighting Biological processes like microtubule organization, DNA endonuclease activity and cell-junction organization.

<u>Cellular components:</u>

**Fig. 6(b):**
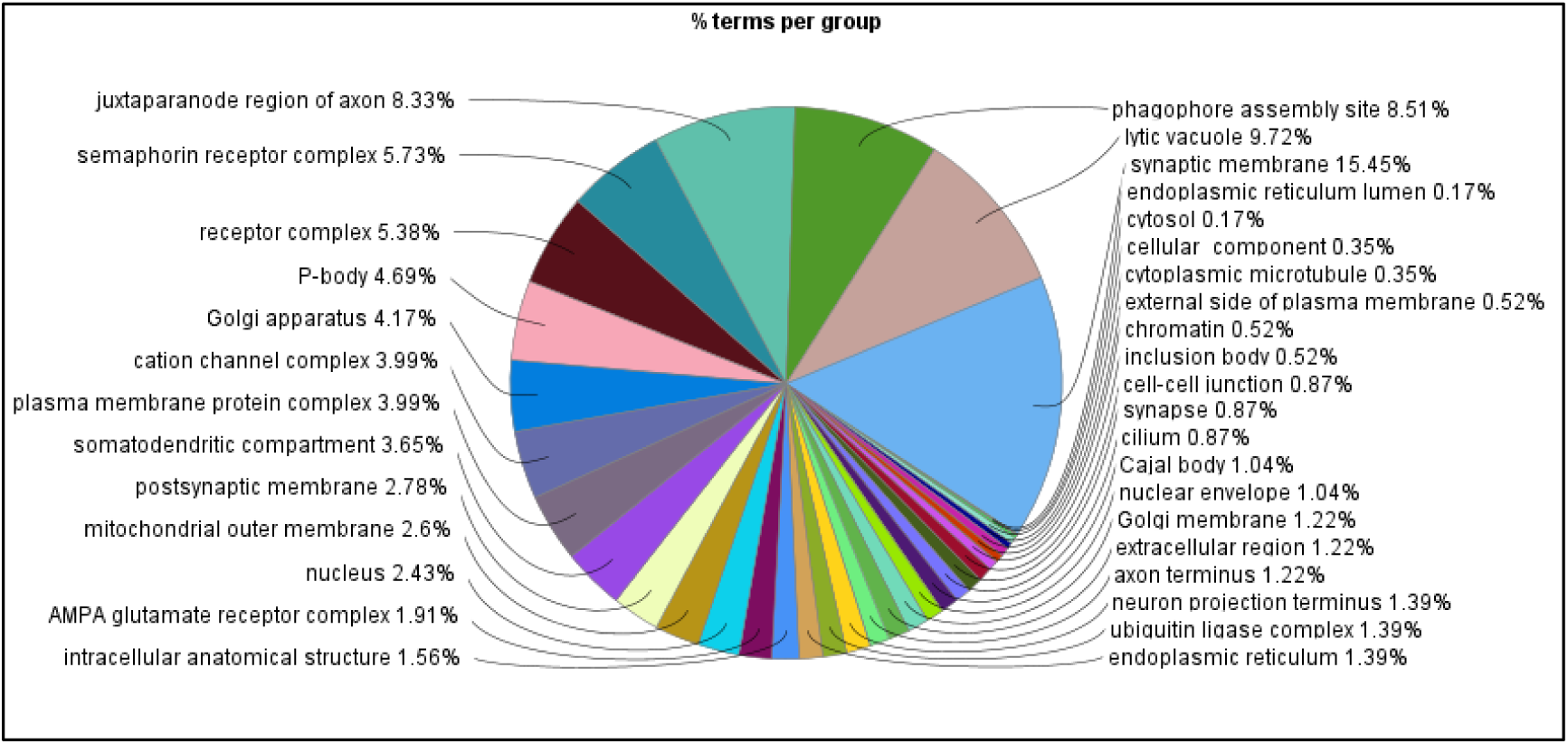
shows pie chart that highlights cellular components like chromatin, nucleus and nuclear envelope.

<u>Molecular functions:</u>

**Fig. 6(c):**
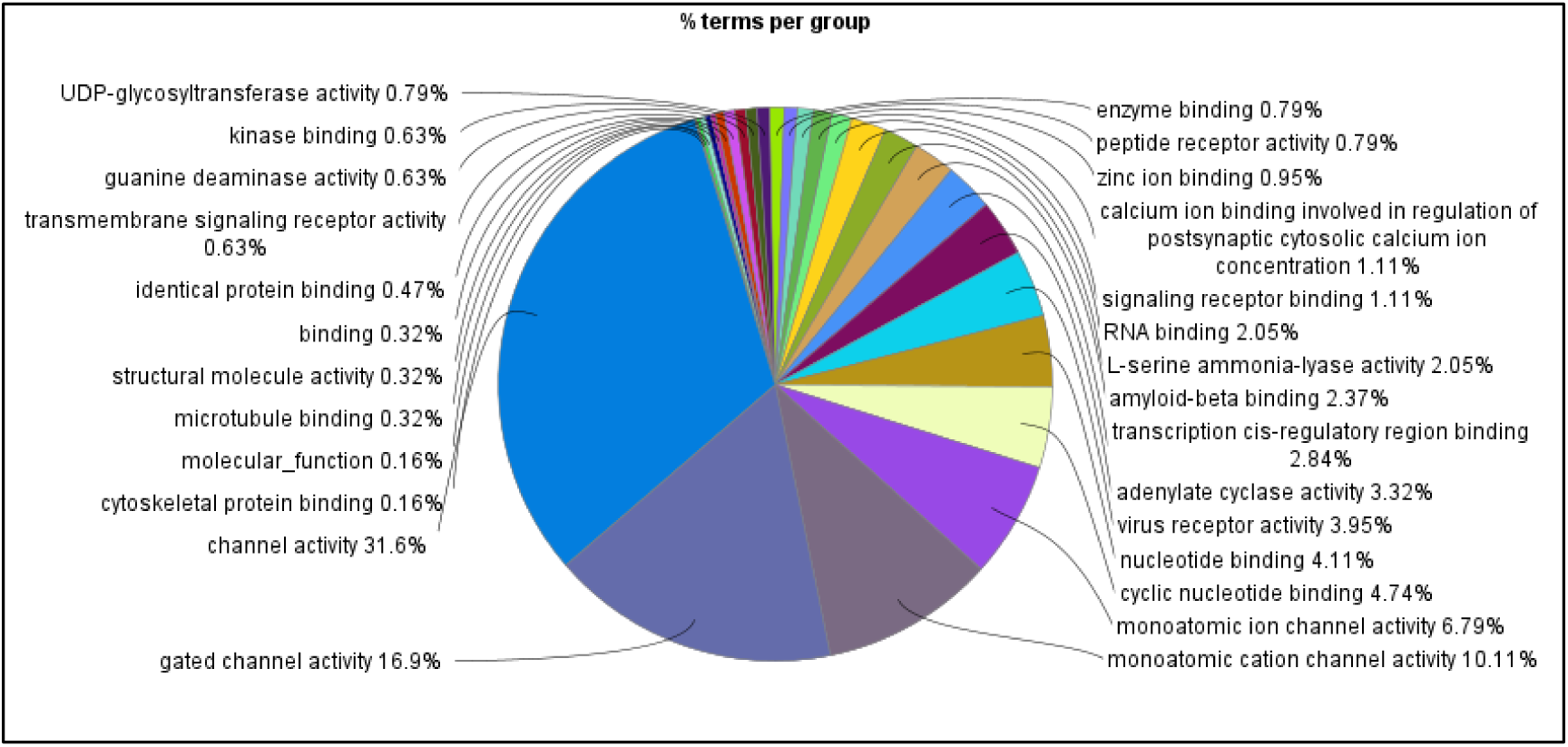
shows pie chart highlighting molecular functions like microtubule binding, cytoskeleton protein binding.

<u>Reactome pathways:</u>

**Fig. 6(d):**
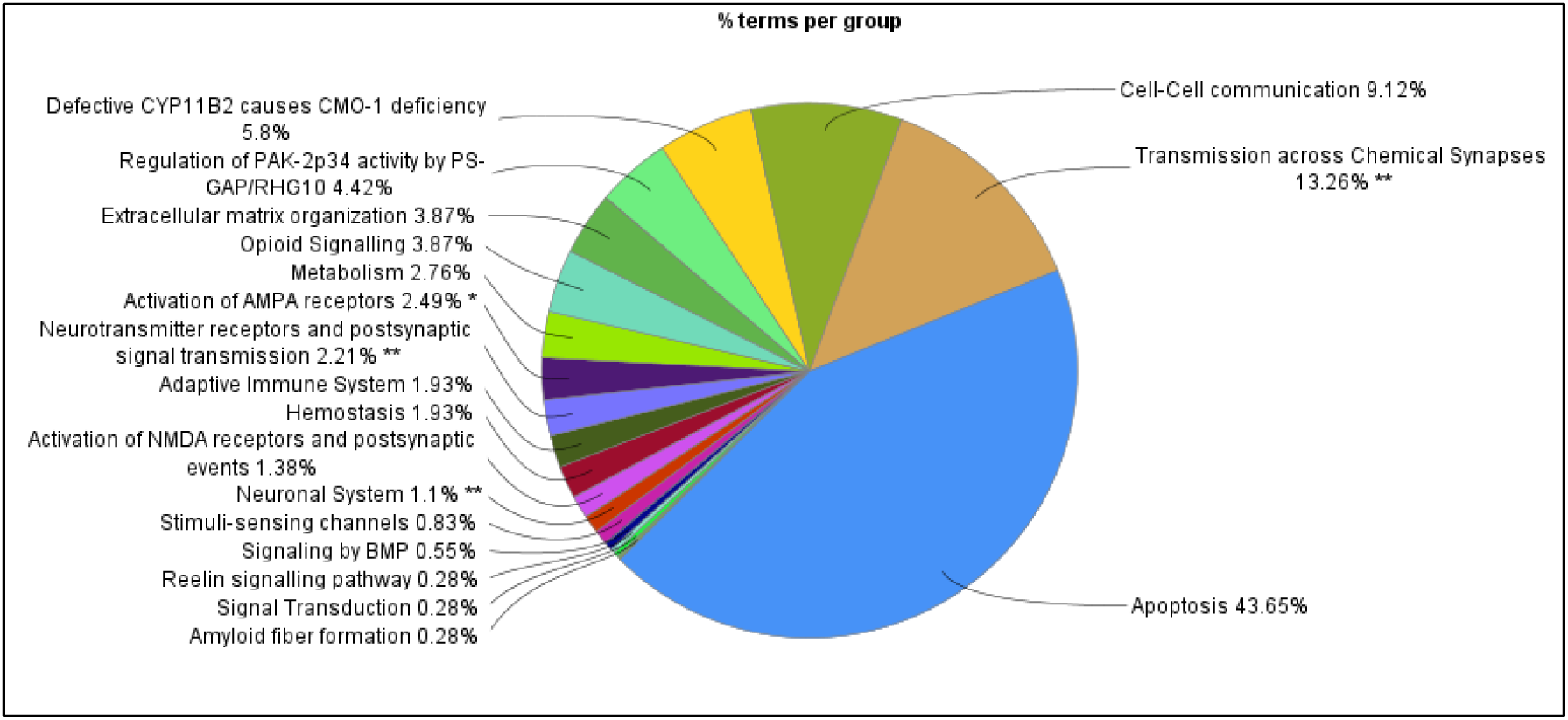
shows pie chart that highlights pathways like apoptosis (for non-repairable cells) and extracellular matrix organization (repairable cells).

**Stage 1 Upregulated & Stage 4 Downregulated genes-**

These genes were found to be aiding tumor progression.

<u>Biological processes:</u>

**Fig. 6(e):**
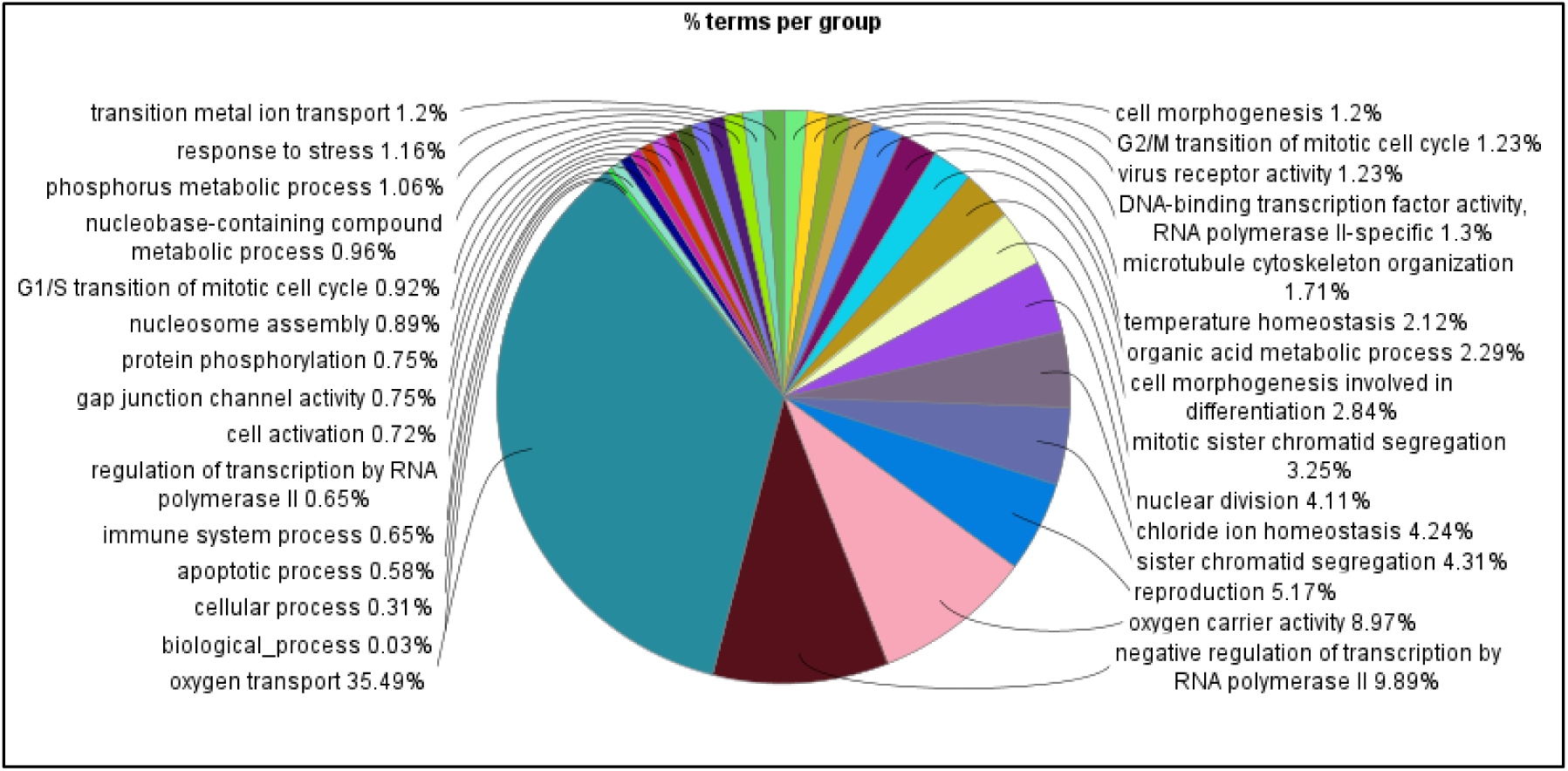
shows pie chart that highlight Biological processes like oxygen transport (important factor for tumor progression) and other cell division processes.

Further analysis associated with these DEGs is enlisted in the Supplementary materials (S4-S11)

Moreover for Melanoma, a set of 17 genes have been shortlisted falling into the TYPE-II criteria (as mentioned in our earlier work **(16)**), which contains those genes that have contrasting or opposite expression values in time-points t1 and t4.

Further, we investigated common factors falling under TYPE-II category in Melanoma and Neuroblastoma and finally found 2 genes common between Melanoma and Neuroblastoma, which are listed in Table 2.

**Table 2:**
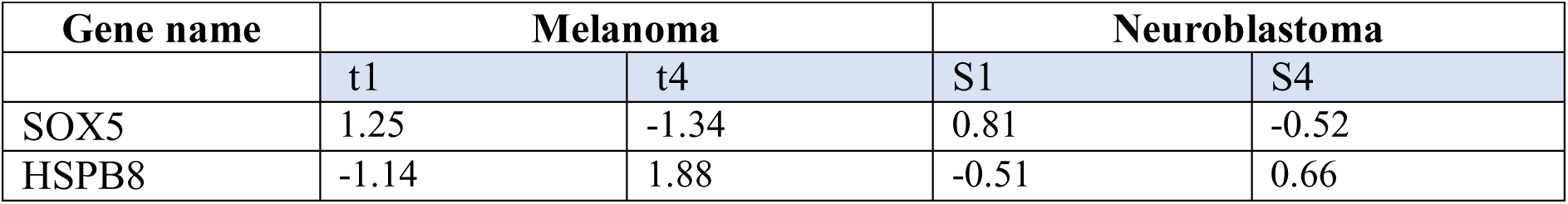
logFc values of the common TYPE-II genes in Melanoma and Neuroblastoma.

The SOX5 gene functions as an oncogene in most of the cancers, promoting cellular proliferation, metastasis, invasion and therapy resistance by stimulating processes like Epithelial-Mesenchymal Transition (EMT). It is shown to be upregulated in progressing phases and downregulated in regressing phases, thereby supporting its oncogenic phenotype. On the contrary, HSPB8 suppresses tumors by promoting cell-death (apoptosis), thereby showing upregulation in tumor regressing stage 4 and time-point t4 in Neuroblastoma and Melanoma respectively. This also supports our hypothesis of leading the unrepairable cells into apoptosis to eliminate tumor cells permanently, as discussed in Section 4.3 (Significance of the study).

### 3.3 Identification of Protagonist gene-factor playing crucial role in tumor regression

Finally, our protagonist gene-SYNE1 marks analysis Type III (Fig. 7), which is also found in Type I as well as Type II gene set as upregulated gene in Stage 4 (Table 3).

**Fig. 7:**
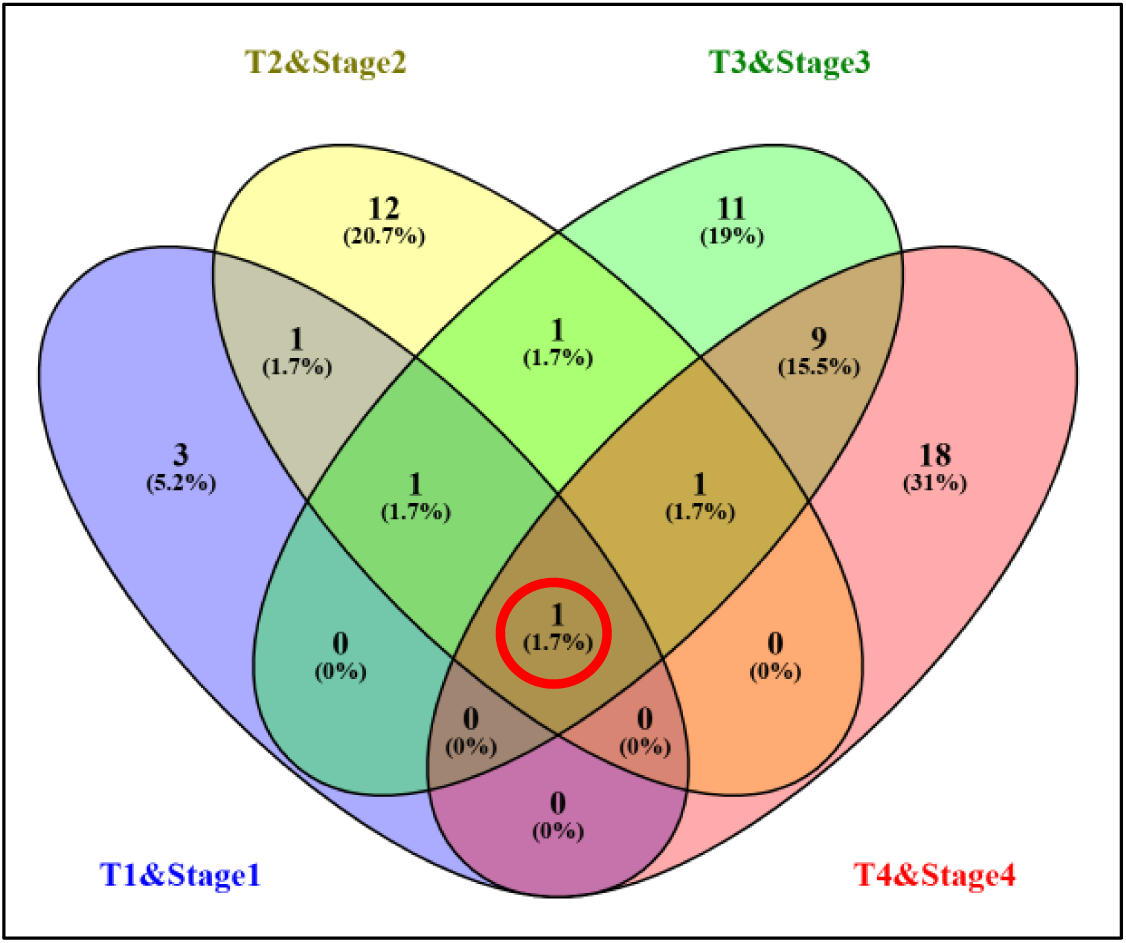
Venn diagram representing the protagonist gene SYNE-1 common in all temporal phases of Melanoma & Neuroblastoma.

**Table 3:** Numerating logFc values of SYNE1 in Melanoma and Neuroblastoma.

| Melanoma |  |  |  | Neuroblastoma |  |  |  |
| --- | --- | --- | --- | --- | --- | --- | --- |
| t1 | t2 | t3 | t4 | S1 | S2 | S3 | S4 |
| 1.03 | 1.11 | 2.47 | 2.07 | -1.17 | -0.16 | 0.62 | 0.59 |

To explore the importance of SYNE1, we constructed a heatmap of its expression in different tissues of the body (Fig. 8).

**Fig. 8:**
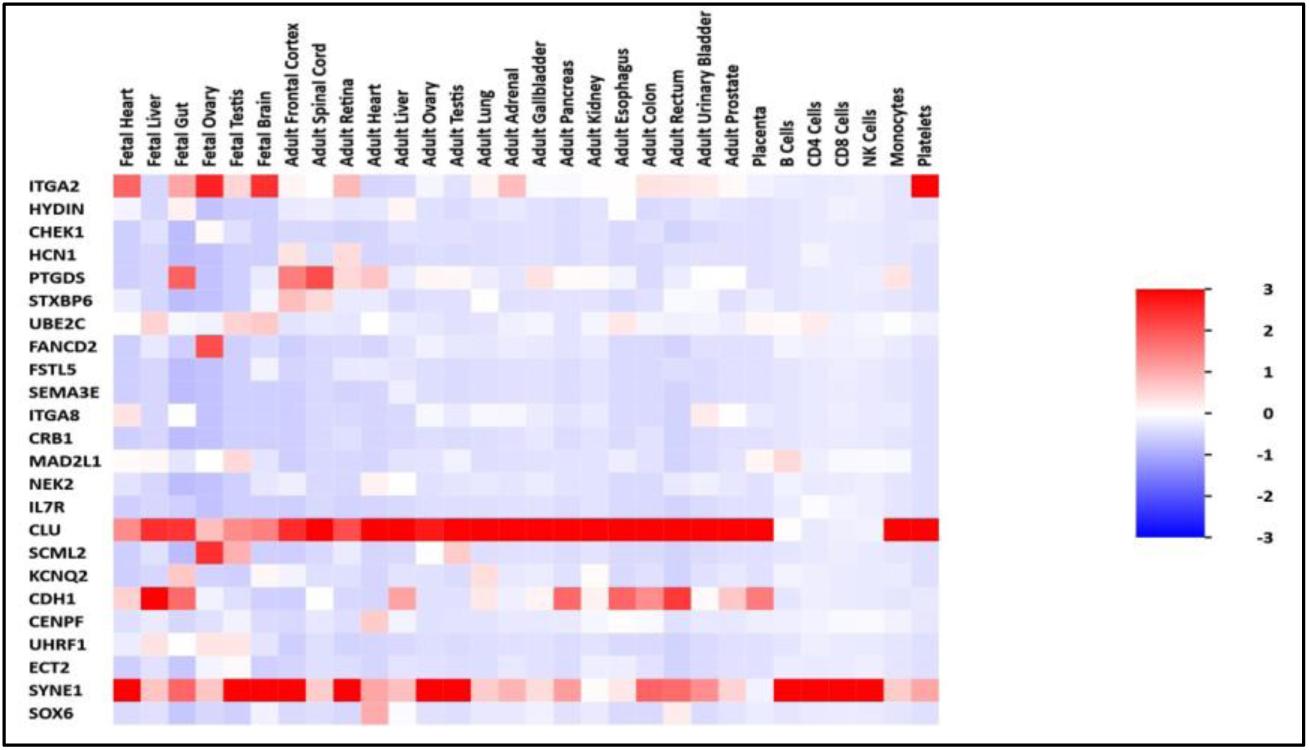
SYNE1 expression level in various tissues.

This analysis by three approaches co-insides with each other, pointing out the commonality and at the same time importance of it which helps us understand the in-depth landscape of pathways that are activated by our system to combat tumor cells thereby leading to tumor regression. These results showed that the upregulated Nesprin-1 was closely related to the DNA damage repair via mis-match repair pathway, which indicates that the deregulation of Nesprin-1 caused by mutation might lead to poor overall survival. Therefore, we also delineated the genetic alteration/ mutation occurring in SYNE-1 using cBioPortal (TARGET-NBL, GDC) which is denoted as VUS (Variant of Uncertain Significance) which means that there is not enough information available about this mutation and therefore it is important to be explored and evaluated (Fig. 9).

**Fig. 9:**
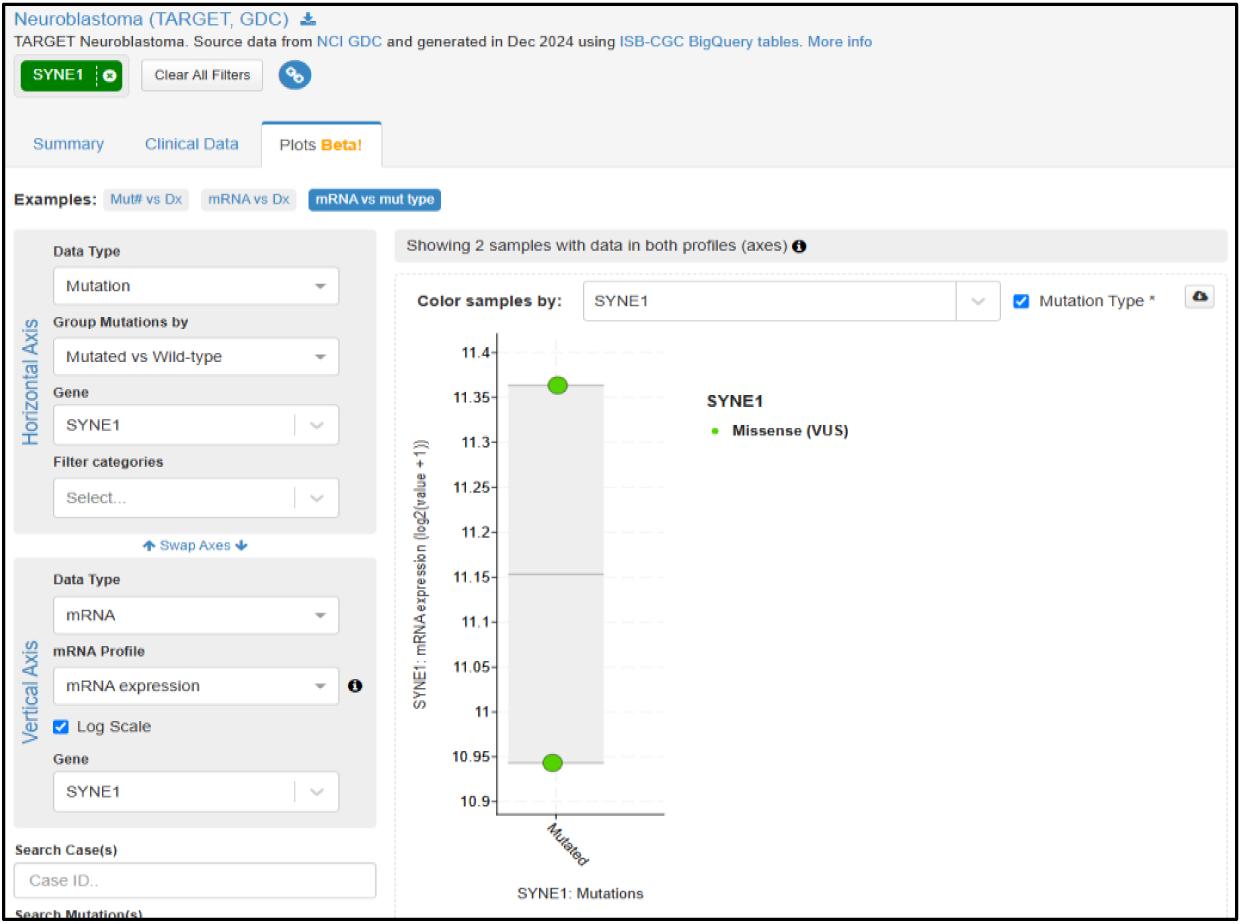
Mutational analysis of SYNE1.

Thereafter, we present the network constructed by the SYNE-1 and its most closely interacting neighboring genes. We shortlisted these neighboring interacting genes viz, SUN1/2, Lamins, Emerin, Actin which might be the novel targets in the future studies (Fig. 10).

**Fig.10:**
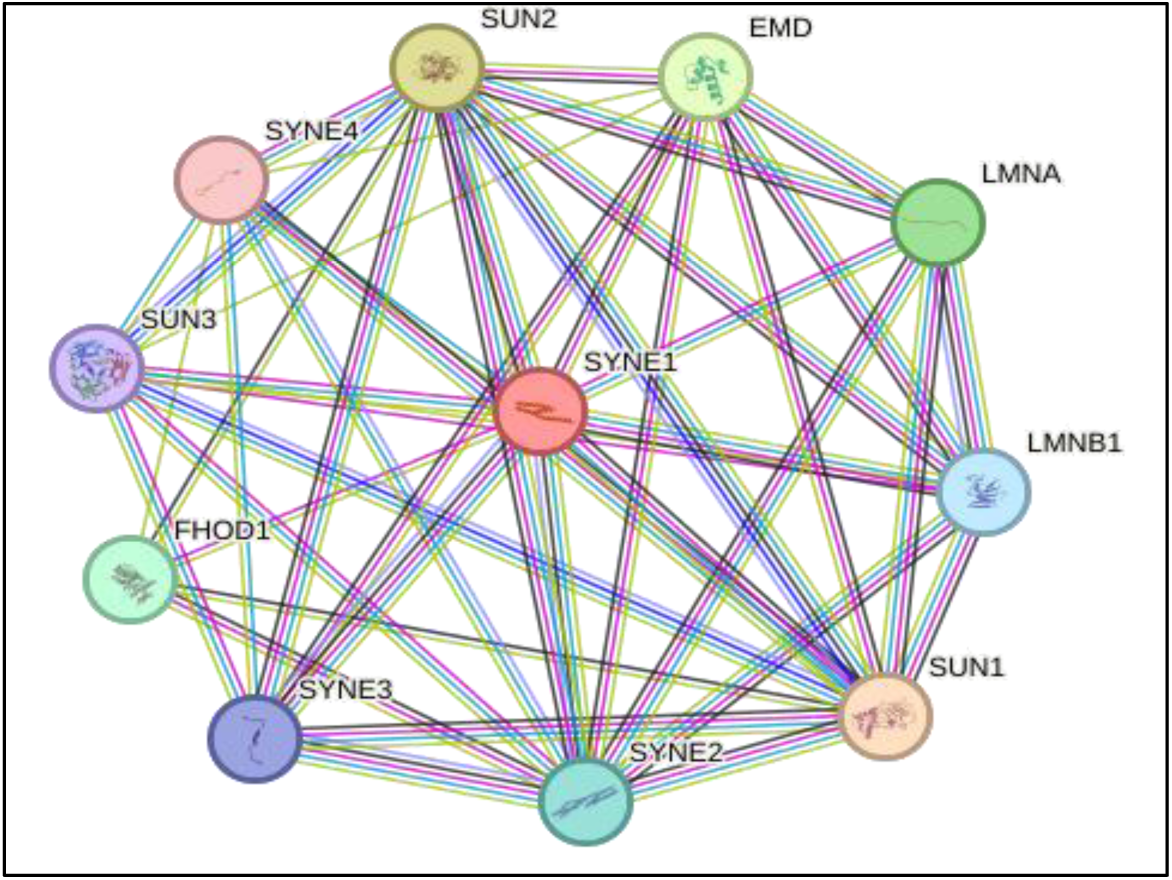
Depicts String network of SYNE1 with its interacting genes showing 11 nodes and 45 edges.

### 3.4 miRNA-protagonist gene network

We also investigated the miRNAs interacting with Nesprin-1 (SYNE1) and found two prominent micro RNAs viz; mir-218-5p and miR-22-3p with the help of NetworkAnalyst (tool to find miRNA interactome of specific mRNA/protein) (https://www.networkanalyst.ca/NetworkAnalyst/Secure/vis/NetworkView.xhtml) (Fig.11). Two miRNAs highlighted in the image were found common in three micro-RNA databases namely-miRViz v1.3, miRDb, and TargetScan 7.2 respectively.

**Fig.11:**
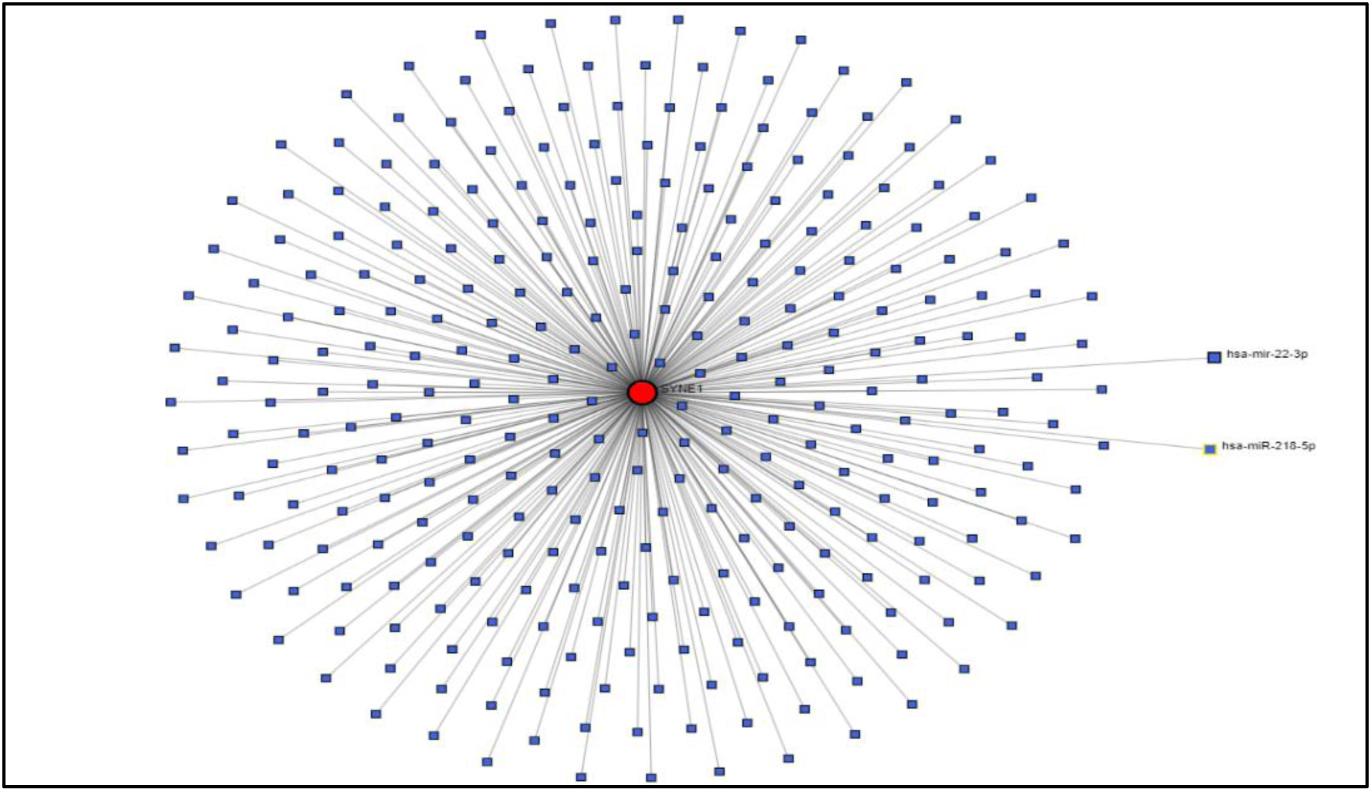
Represents miRNA interactome of Nesprin-1, highlighting miR-22-3p and miR-218-5p.

We then investigated miRNA network interacting with Nesprin-1 with the help of miRViz v1.3 (Fig. 12b), which redirected us to miRDb confirming the two miRNAs and further at TargetScan.

**Fig 12:**
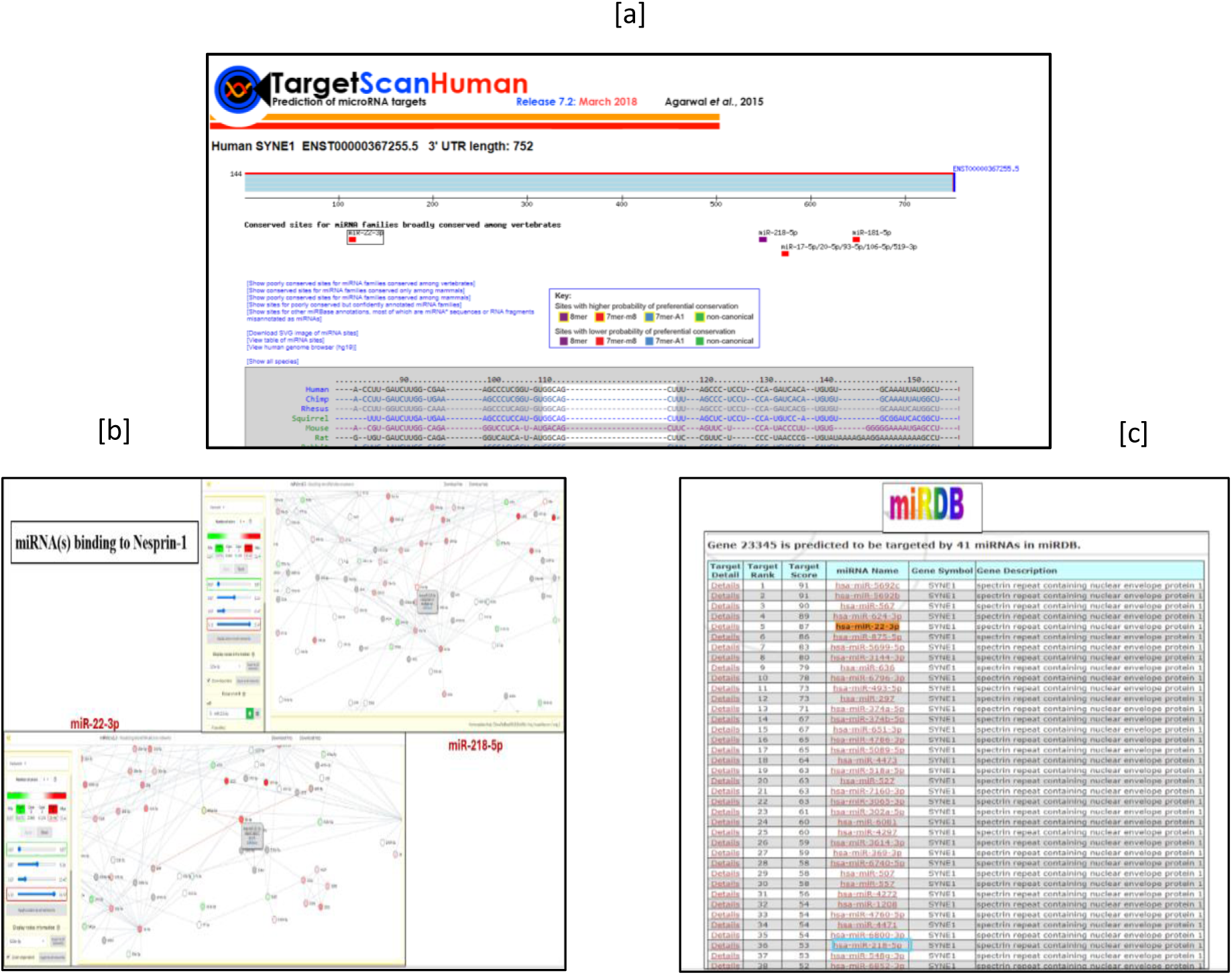
Databases verifying miRNA investigation against SYNE-1 (a)TargetScan 7.2 (b) miRViz v1.3 (c) miRDB.

Moreover, as seen in the images below,-Firstly, miR-22-3p is a 7mer-m8 (Fig. 12a), which means it has an exact match with the seed region + 8^th^ position. Secondly, it has a Target score of 87 as compared to miR-218-5p which has a target score of 53 (Fig. 12c), and as per database guidelines (https://mirdb.org/faq.html), score above 80 is considered to be good and confident, so we went ahead with miR-22-3p and explored it’s interaction with SYNE-1 gene, so as to find out their binding strength to the gene.

Henceforth, we investigated the role of miR-22-3p in various tumors by reviewing the literature available (Fig. 13).

**Fig. 13:**
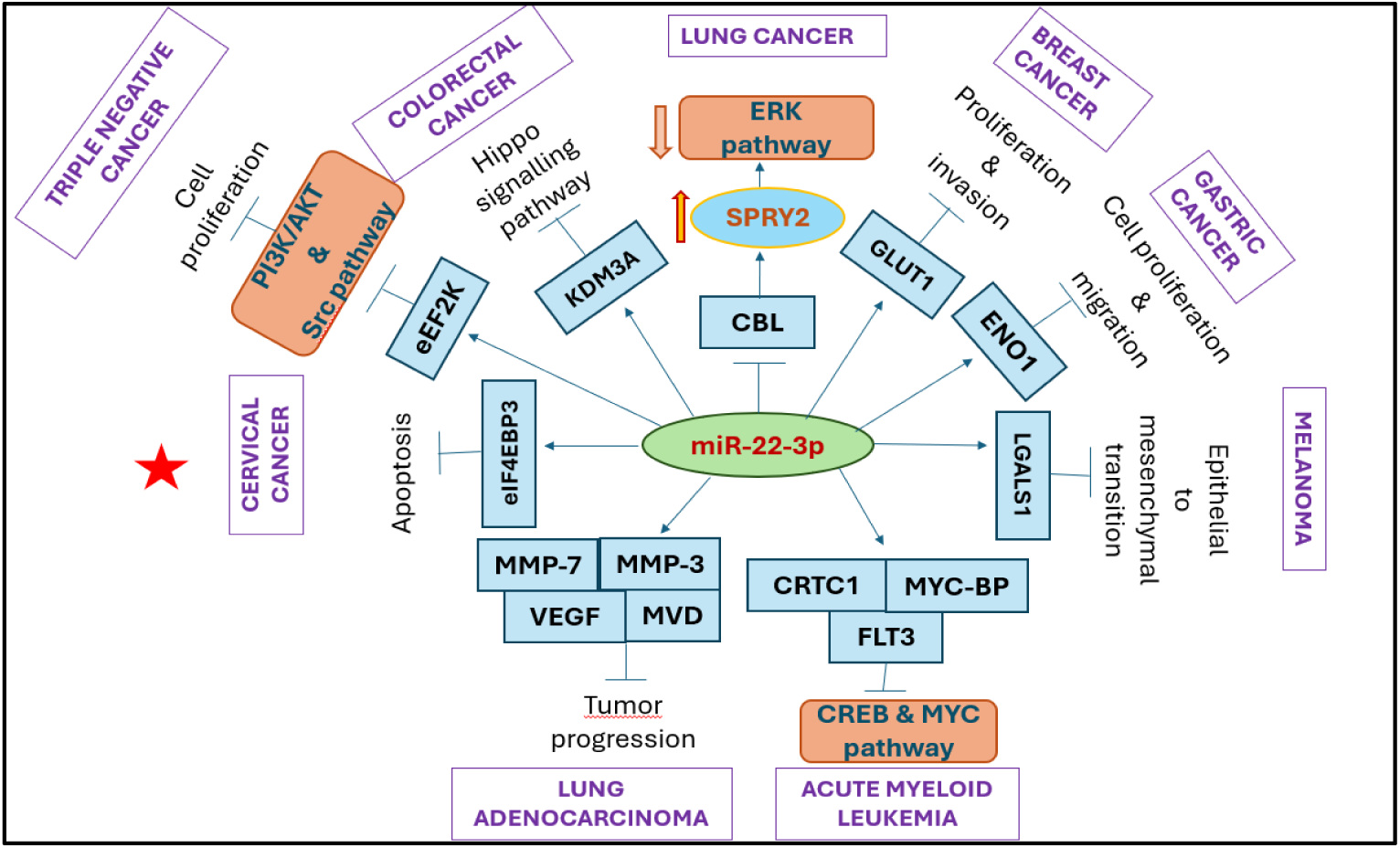
represents role of miR-22-3p in various tumors.

To further verify the importance and binding strength of miR-22-3p (5’AGCUGCC 3’), we analyzed it’s binding using RNA Hybrid bibiserve (https://bibiserv.cebitec.uni-bielefeld.de/rnahybrid/) which demonstrates the maximum free energy (mfe) value generated upon binding.

There are two criteria’s which indicate strong binding-

i. The mfe value should be around −30Kcal/mol
ii. The miRNA seed region should be almost entirely complimentary to the mRNA binding region.

So, as seen in Figure 14, upon binding of miR-22-3p the mfe generated is −30Kcal/mol as compared to miR-218-5p which generates an mfe of −25.3 Kcal/mol, we choose to further analyze its interacting mechanism and investigate the pathways related to miR-22-3p and Nesprin-1 for our study.

**Fig. 14:**
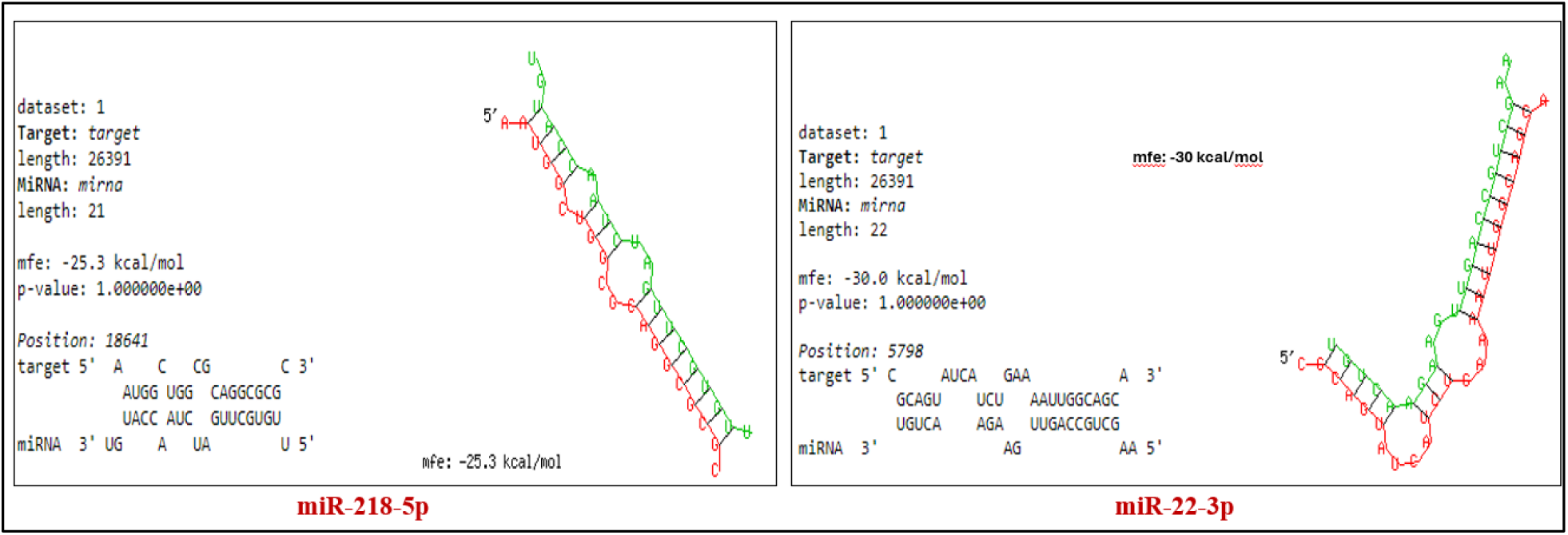
demonstrates the binding strength of miR-218-5p and miR-22-3p against SYNE-1.

### 3.5 Spontaneous tumor regressing Pathway analysis

Once we found the interacting proteins around SYNE-1 (Nesprin-1), we delineated the path followed by Nesprin-1 for repairing the DNA damage in the cells and the mechanism of repairing the damage.

This investigation led us to understand the involvement of Nesprin-1 in mis-match repair mechanism where Mut-S complex helps in locating the mis-match. Nesprin-1 has been found to play crucial role in the formation of Mut-S complex without which the repairing of cells is not possible. Thus, we uncovered the role of Nesprin-1 in tumor regression via mis-match repair pathway (Fig 15).

**Fig.15:**
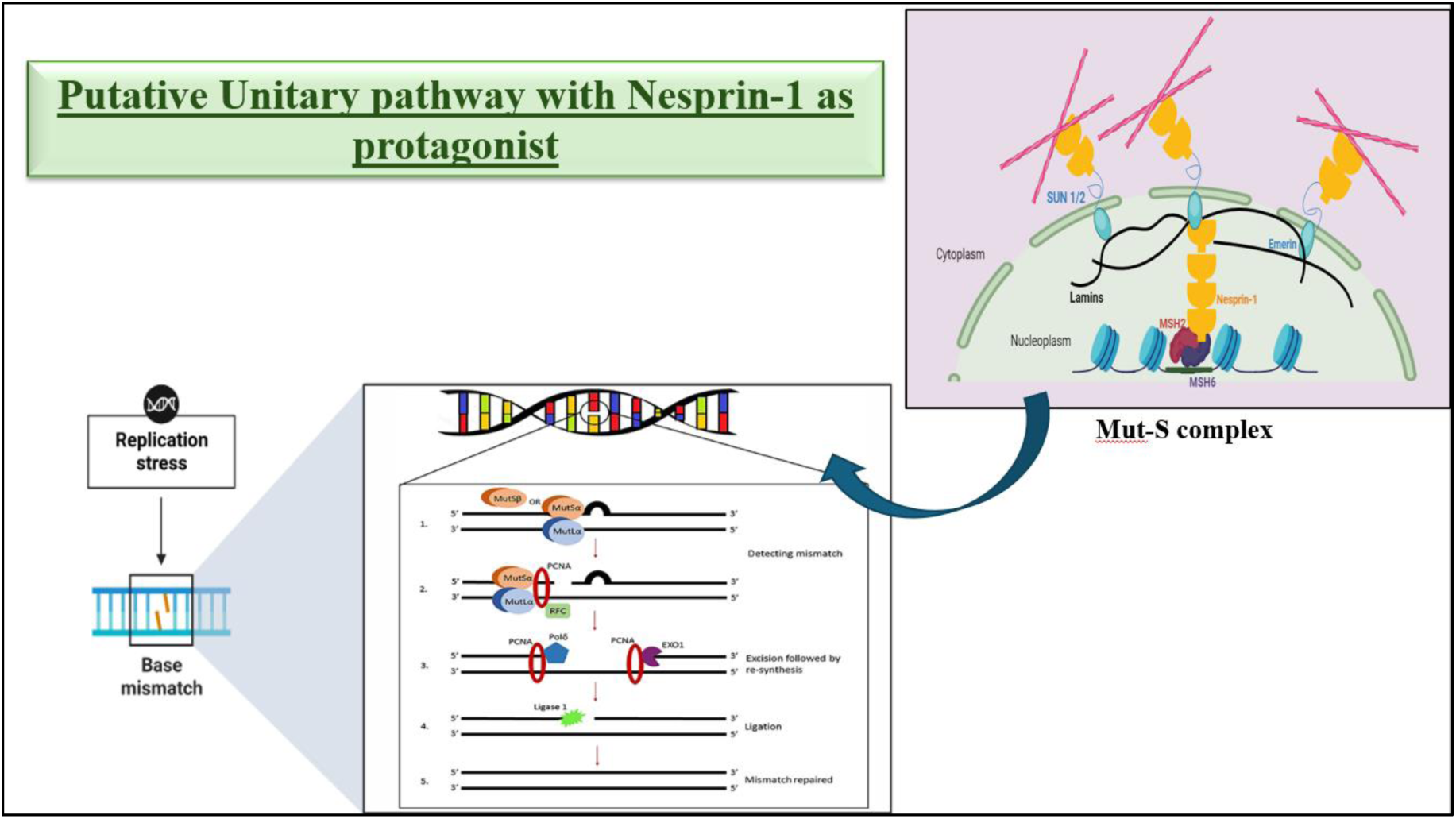
Putative pathway of DNA repair mechanism of malignant cells undergoing tumor regression.

Furthermore, we investigated the pathway followed by SYNE-1 and miR-22-3p to inhibit cellular proliferation and surprisingly found that both the gene as well as miRNA targets PI3K/AKT pathway via respective paths as elaborated in Fig. 16.

**Fig. 16:**
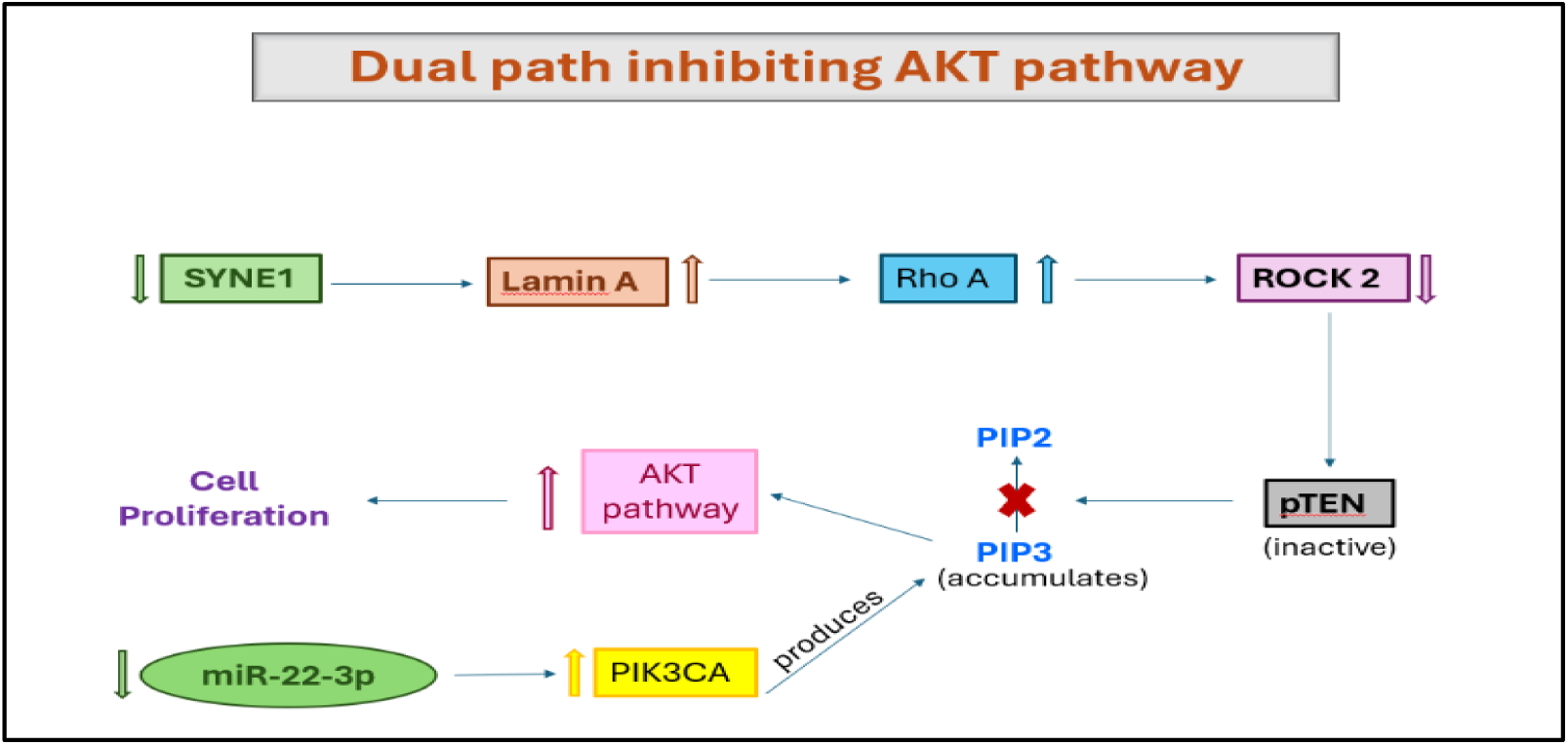
Represents Dual path inhibiting PI3K/AKT.

## 4. DISCUSSION

Spontaneous tumor regression, a rare spontaneous phenomenon where tumor shrinks and disappears without any kind of conventional treatment. This astonishing event needs to be studied thoroughly to understand its underlying mechanism and genetic landscape contributing to regression or remission of tumor cells.

### 4.1 Associated genes & Pathways

In this study, we have probed the microarray dataset of Neuroblastoma patients conducted on Agilent microarray chip, available at Gene expression Omnibus. This helped us to trace down the most significant genes and their associated biological pathways that putatively act as the driver for tumor regression process. We analyzed the upregulated and downregulated DEGs in each stage of Neuroblastoma and time point of Melanoma (as discussed in our previous work), and categorized our study into three different groups-

Type I, included the genes showing consistent expression across tumor regressing phases of Melanoma-t2, t3, t4 and Neuroblastoma-S3, S4.

Type II, included the genes showing contrasting or opposite expression in progressing phase-t1 and S1 in contrast to t4 and S4 phases of Melanoma and Neuroblastoma respectively.

Type III, included the only gene factor we traced as a common gene expressing consistently across all time points of Melanoma and stages of Neuroblastoma, being true to its nature acting as tumor suppressor.

We performed Enrichment analysis for the genes in each type and discovered that in each type, the genes are associated with cell-junctions organization, extracellular matrix organization, cellular DNA damage repair and apoptosis. The most commonly enriched pathways demarcated by the driver genes are involved in (i) cellular replication, (ii)cell-cycle regulation and division, (iii) DNA-damage repair, (iv) signaling pathways like PI3K/AKT pathway.

### 4.2 Protagonist gene & associated miRNA inducing spontaneous tumor regression

The gene associated with Type III of our investigation is our protagonist genes-SYNE1 which is a tumor suppressor gene, thoroughly studied in cancer progression but its role in tumor regression is yet to be explored. SYNE-1 is a spectrin-repeat containing nuclear envelope protein-1 which connects nucleus with the cytoplasm. It has been extensively studied for an autosomal recessive disorder that is Cerebellar ataxia **[24]**. We also, investigated the protein coded by SYNE1 which is Nesprin-1. Nesprin-1 is a linker protein, that is associated with the nuclear membrane and binds to F-actin through the N-terminus, while via C-terminus it binds to SUN proteins which is a transmembrane protein **[25]**. This interaction is crucial for the formation of LINC complex **[26,27]**. We mined the data available extensively and found that this LINC complex and its interaction with SUN and Lamin proteins is extremely crucial for formation of Mut-S complex, which ultimately leads to repairing of damaged/ mis-matched DNA via Mis-match repair pathway. This, also helped us to demarcate the role of Nesprin-1 in maintaining the shape and cellular integrity of the cell, and that its deficiency (due to mutation) can lead to loss of well-defined nuclear and cytoskeletal architecture, thereby resulting in misshapen nucleus, altered nuclear envelope arrangements and dislodging of centrosome from the nucleus, converting healthy normal cells into distorted tumor cells **[28]**. Thereafter, we precisely constructed an interactome of SYNE1 gene, including the genes functionally related to it as well as the microRNAs that target this gene. We found that the miRNA with accurate seed-region matching, and maximum interaction score with SYNE1, as analyzed through various non-coding databases is miR-22-3p. Thus, we further explored the common pathways targeted by SYNE1 and miR-22-3p and found that they both target PI3K/Akt pathway following different paths and mechanisms (Fig. 17).

**Fig. 17:**
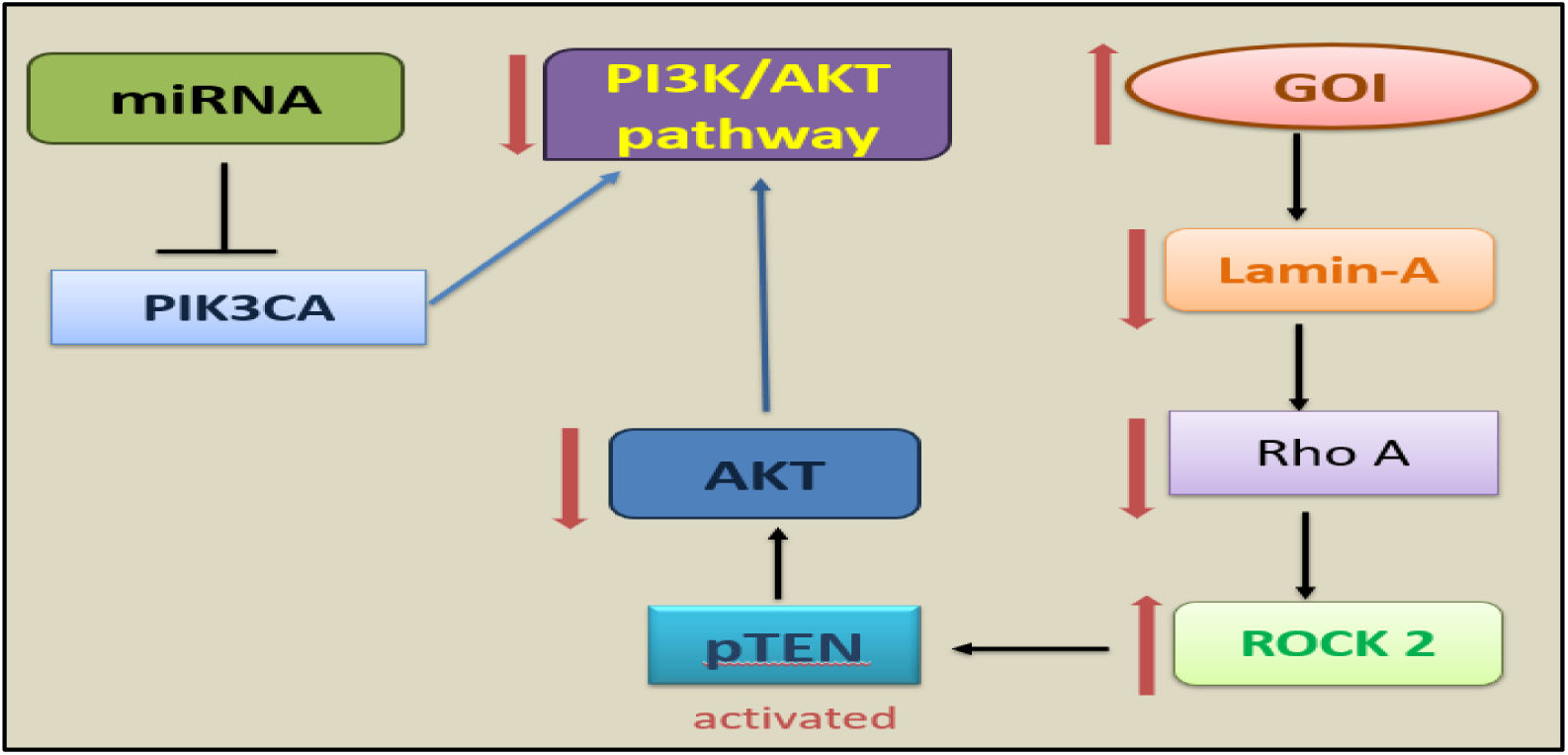
Represents connecting path between miR-22-3p and SYNE-1.

### 4.3 Significance and Implications of Study

PI3K/Akt pathway is a well-known pathway that is over-activated in several human cancers, and leads to cellular proliferation, invasion and metastasis of tumor cells thereby contributing to carcinogenesis **[29]**. In our study, we have found that our protagonist gene SYNE1 as well as the miRNA (miR-22-3p) targeting it inhibits this pathway thereby helping the tumor cells to undergo apoptosis. Simultaneously, we have also found that our protagonist gene SYNE1 codes for Nesprin-1 which is involved in mis-match repair pathway, thereby aiding DNA repair of the damaged tumor cells as well as maintaining the cellular structure and integrity.

Thus, with the help of this study, we can conclude that SYNE1 might be playing a crucial role in tumor regression phenomenon by two mechanisms-

i. Reverting DNA-damage in the tumor cells that can be converted into healthy cells, using mis-match repair mechanism involving Nesprin-1, and
ii. Leading unrepairable tumor cells into apoptosis by inhibiting PI3K/Akt pathway via-two different paths involving miR-22-3p.

## 5. CONCLUSION

This study formulates a new functional aspect of sparsely investigated protein, involved in DNA repair mechanisms during spontaneous regression of malignant tumors. We found in this study, role of Nesprin-1 protein in its true nature as a tumor suppressor in Melanoma as well as Neuroblastoma and because of our study we can conclude that this protein Nesprin-1 may have relevance in SCR (Spontaneous cancer regression) of other malignant tumors.

Since we found that the miRNA (miR-22-3p), found interacting with SYNE-1 gene itself targets PI3K pathway enabling SCR, we can say that the malignant cells undergoing SCR are either going through the repair mechanism involving Nesprin-1 (malignant cells which can be repaired) or Apoptosis through PI3K pathway (malignant cells which cannot be repaired). Thus, we need to target these two paths through which we can formulate therapeutics which can aid in SCR.

**Supplementary Materials** for this work are available at the end of this document.

## Notes

### Competing Interest Statement

The authors have declared no competing interest.

## REFERENCES

1. Zahl PH, Gøtzsche PC, Mæhlen J. Natural history of breast cancers detected in the Swedish mammography screening programme: a cohort study. Lancet Oncol. 2011 Nov;12(12):1118–24.

2. Fryback DG, Stout NK, Rosenberg MA, Trentham-Dietz A, Kuruchittham V, Remington PL. The Wisconsin Breast Cancer Epidemiology Simulation Model. J Natl Cancer Inst Monogr. 2006;(36):37–47.

3. PDQ Pediatric Treatment Editorial Board. Neuroblastoma Treatment (PDQ(R)): Health Professional Version. In: PDQ Cancer Information Summaries. Bethesda (MD): National Cancer Institute (US); 2002.

4. Gurney JG, Ross JA, Wall DA, Bleyer WA, Severson RK, Robison LL. Infant cancer in the U.S.: histology-specific incidence and trends, 1973 to 1992. J Pediatr Hematol Oncol. 1997; 19(5): 428–432.

5. London WB, Castleberry RP, Matthay KK, et al. Evidence for an age cutoff greater than 365 days for neuroblastoma risk group stratification in the Children’s Oncology Group. J Clin Oncol. 2005; 23(27): 6459–6465.

6. Maris JM. Recent advances in neuroblastoma. N Engl J Med. 2010; 362(23): 2202–2211.

7. Siegel RL, Miller KD, Wagle NS, Jemal A. Cancer statistics, 2023. CA Cancer J Clin. 2023 Jan;73(1):17-48.

8. Wong VK, Lubner MG, Menias CO, Mellnick VM, Kennedy TA, Bhalla S, Pickhardt PJ. Clinical and Imaging Features of Noncutaneous Melanoma. AJR Am J Roentgenol. 2017 May;208(5):942–959.

9. Eisen MB, Brown PO. DNA arrays for analysis of gene expression. Methods Enzymol. 1999;303:179–205.

10. van de Vijver MJ. The role of gene expression profiling by microarray analysis for prognostic classification of breast cancer. Breast Cancer Res. 2005;7(Suppl 1):S1.

11. Wang, C., et al. Assessment of genetic testing and related counseling services: Current research and future directions. Social Science and Medicine 58, 1427—1442 (2004)

12. Bittner M, Meltzer P, Chen Y, Jiang Y, Seftor E, Hendrix M, Radmacher M, Simon R, Yakhini Z, Ben-Dor A, Sampas N, Dougherty E, Wang E, Marincola F, Gooden C, Lueders J, Glatfelter A, Pollock P, Carpten J, Gillanders E, Leja D, Dietrich K, Beaudry C, Berens M, Alberts D, Sondak V. Molecular classification of cutaneous malignant melanoma by gene expression profiling. Nature. 2000;406:536–540.

13. Mariadason JM, Augenlicht LH, Arango D. Microarray analysis in the clinical management of cancer. Hematol Oncol Clin North Am. 2003 Apr;17(2):377–87.

14. Al Moustafa, AE., Alaoui-Jamali, M., Batist, G. et al. Identification of genes associated with head and neck carcinogenesis by cDNA microarray comparison between matched primary normal epithelial and squamous carcinoma cells. Oncogene 21, 2634–2640 (2002).

15. Sur-Erdem I, Hussain MS, Asif M, Pınarbası N, Aksu AC, Noegel AA. Nesprin-1 impact on tumorigenic cell phenotypes. Mol Biol Rep. 2020 Feb;47(2):921–934.

16. Kumari B, Sakode C, Lakshminarayanan R, Purohit P, Bhattacharjee A, Roy PK. A mechanistic analysis of spontaneous cancer remission phenomenon: identification of genomic basis and effector biomolecules for therapeutic applicability. 3 Biotech. 2023 Apr;13(4):113.

17. Szklarczyk D, Gable AL, Lyon D, et al. (2019) STRING v11: Protein–protein association networks with increased coverage, supporting functional discovery in genome-wide experimental datasets. Nucleic Acids Res 47: D607–D613.

18. Bindea G, Mlecnik B, Hackl H, et al. (2009) ClueGO: a Cytoscape plug-in to decipher functionally grouped gene ontology and pathway annotation networks. Bioinformatics 25:1091–1093.

19. Pathan M, Keerthikumar S, Ang C-S, et al. (2015) FunRich: An open access standalone functional enrichment and interaction network analysis tool. Proteomics 15: 2597–2601.

20. Dennis G, Sherman BT, Hosack DA, et al. (2003) DAVID: Database for Annotation, Visualization, and Integrated Discovery. Genome Biology 4: R60.

21. Palmieri G, Ombra M, Colombino M, et al. (2015) Multiple Molecular Pathways in Melanomagenesis: Characterization of Therapeutic Targets. Front Oncol 5: 183.

22. Lo JA, Fisher DE (2014) The melanoma revolution: from UV carcinogenesis to a new era in therapeutics. Science 346: 945–949.

23. Benchia D, Bîcă OD, Sârbu I, Savu B, Farcaș D, Miron I, Postolache AL, Cojocaru E, Abbo O, Ciongradi CI. Targeting Pathways in Neuroblastoma: Advances in Treatment Strategies and Clinical Outcomes. Int J Mol Sci. 2025 May 15;26(10):4722.

24. Beaudin M, Gamache PL, Gros-Louis F, et al. SYNE1 Deficiency. 2007 Feb 23.

25. Chancellor TJ, Lee J, Thodeti CK, Lele T. Actomyosin tension exerted on the nucleus through nesprin-1 connections influences endothelial cell adhesion, migration, and cyclic strain-induced reorientation. Biophys J. 2010 Jul 7;99(1):115–23.

26. Starr, D.A., and M. Han. 2002. Role of ANC-1 in tethering nuclei to the actin cyctoskeleton. Science. 298:406–409.

27. Zhang, Q., J.N. Skepper,…, C.M. Shanahan. 2001. Nesprins: a novel family of spectrin-repeat-containing proteins that localize to nuclear membrane in multiple tissues. J. Cell Sci. 114:4485–4498.

28. Sur-Erdem I, Hussain MS, Asif M, Pınarbası N, Aksu AC, Noegel AA. Nesprin-1 impact on tumorigenic cell phenotypes. Mol Biol Rep. 2020 Feb;47(2):921–934.

29. Rascio F, Spadaccino F, Rocchetti MT, Castellano G, Stallone G, Netti GS, Ranieri E. The Pathogenic Role of PI3K/AKT Pathway in Cancer Onset and Drug Resistance: An Updated Review. Cancers (Basel). 2021 Aug 5;13(16):3949.

