## Supplementary materials for "Deciphering a common code: Unitary gene profile factor enabling Spontaneous Tumor Regression"

**3.1 Microarray data analysis and identification of differentially expressed gene**

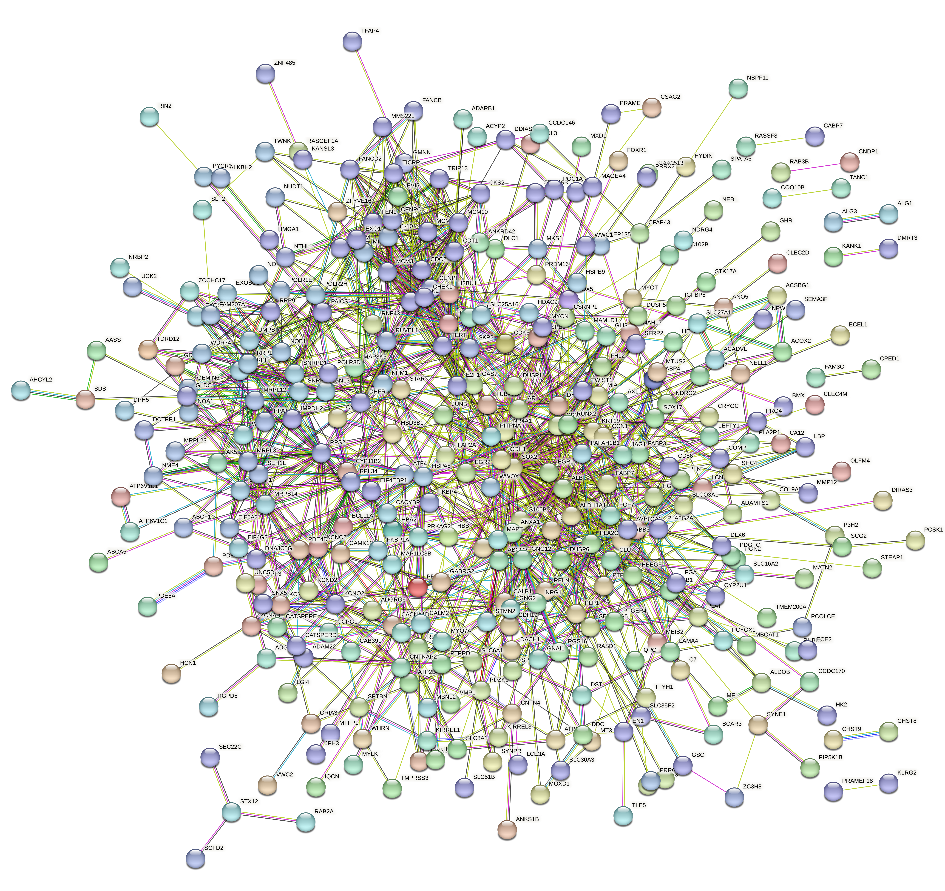

Fig. S1: STRING network depicting connection between 261 DEGs with 493 nodes and 1256 edges during different stages of Tumor regression

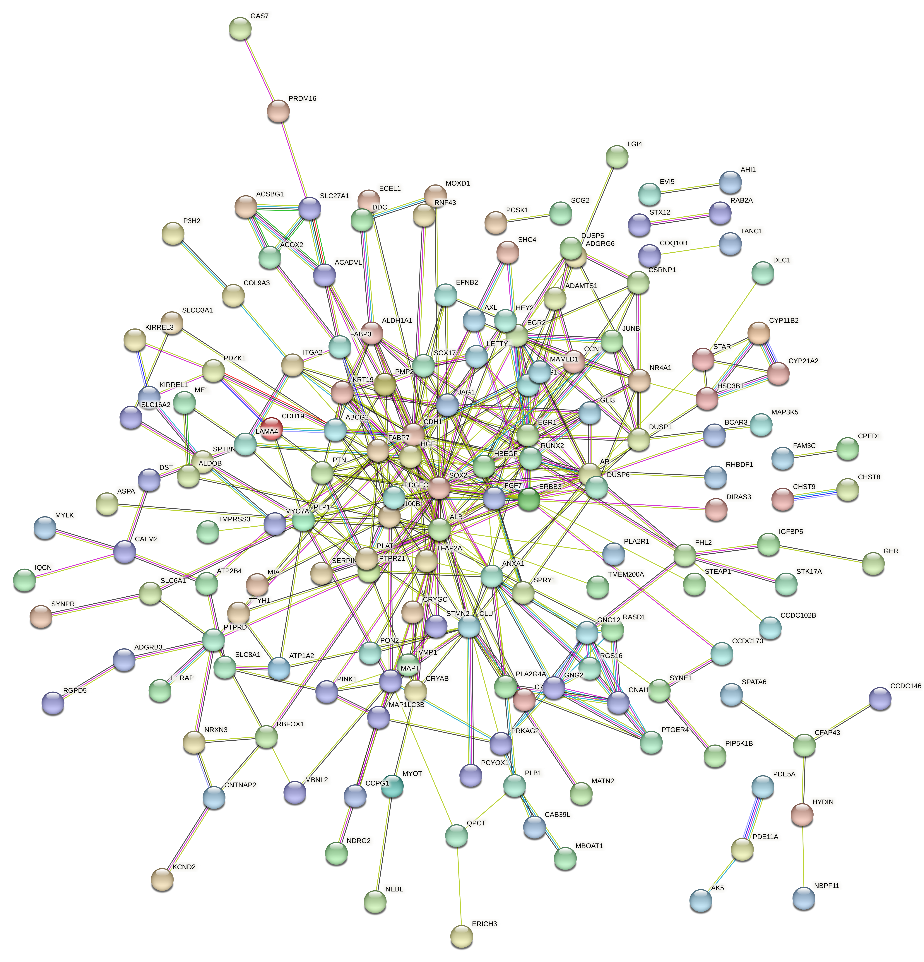

Fig. S2: STRING network showing Upregulated DEGs in stages 3 & 4 in Neuroblastoma, with 258 nodes and 371 edges

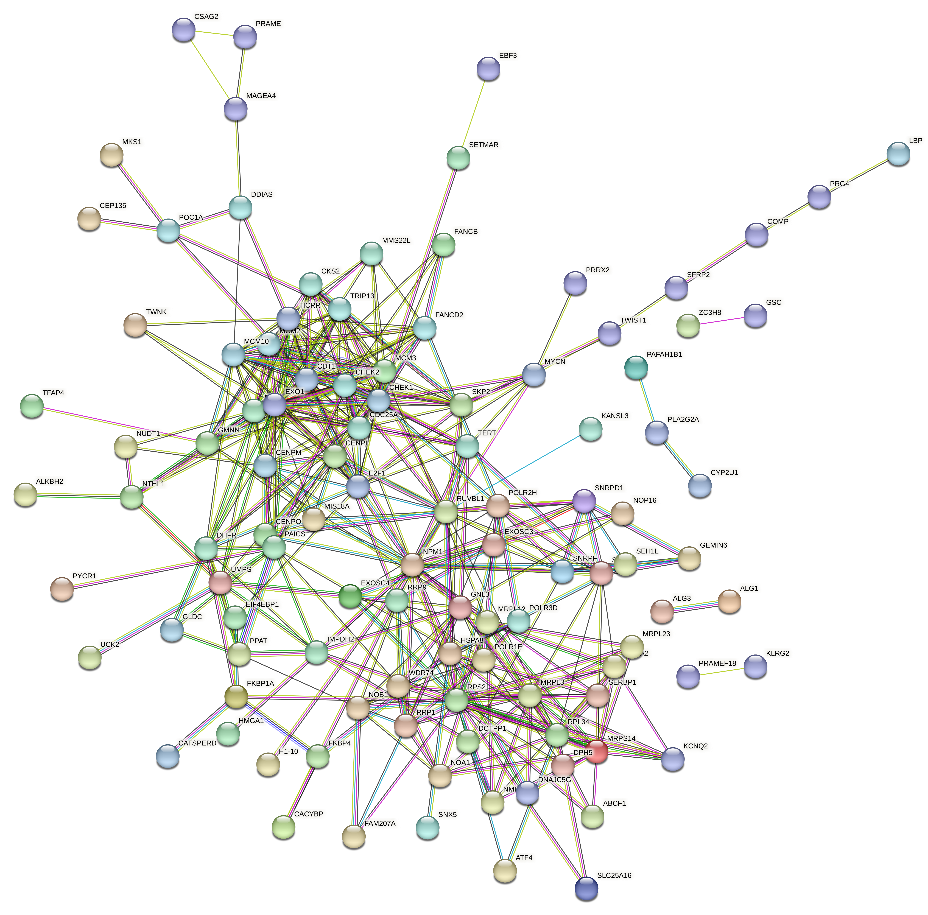

Fig. S3: STRING network depicting Downregulated DEGs in stage 3 & 4 of Neuroblastoma, with 161 nodes and 392 edges

3.2 **Identification of Protagonist gene-factor playing crucial role in tumor regression**

Biological process:

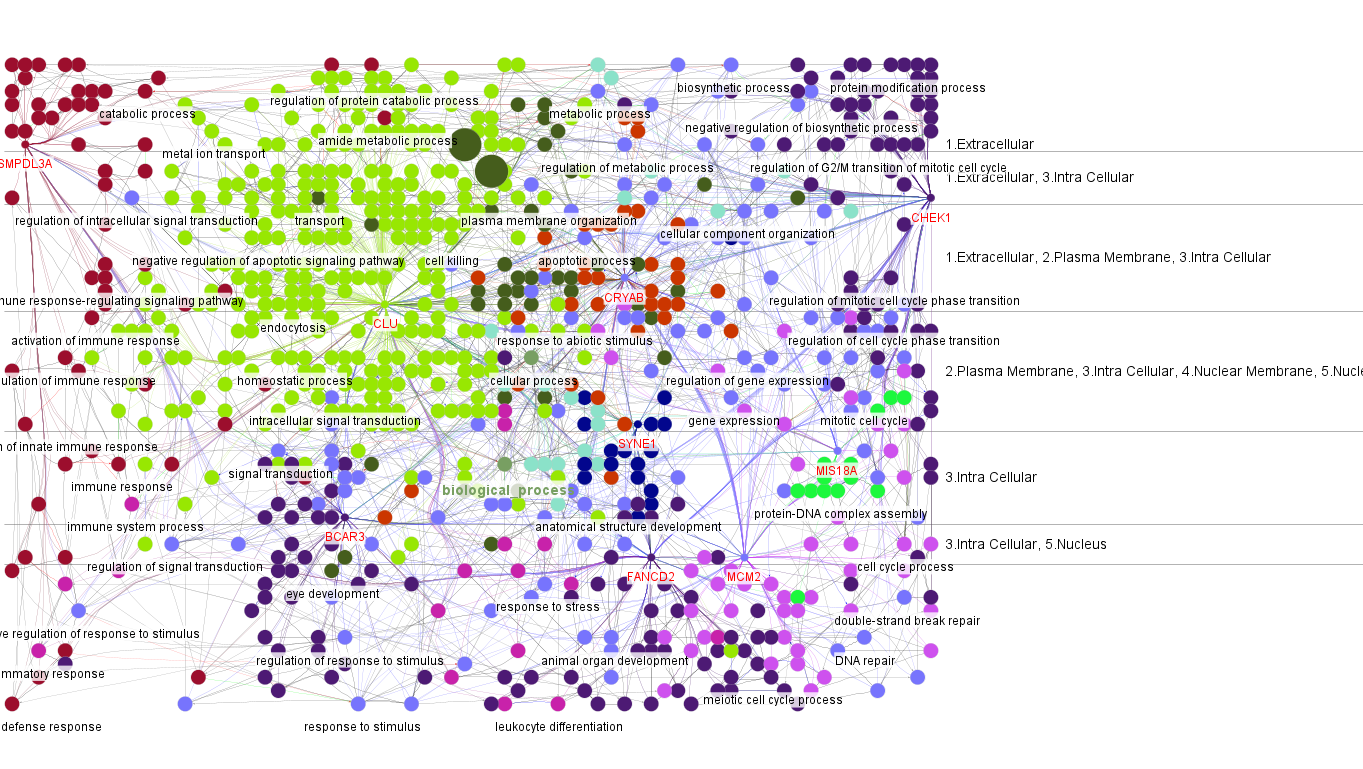

Fig. S4: Cerebral layout of the DEGs highlighting biological processes like nuclear organization and nuclear matrix anchoring

Molecular functions-

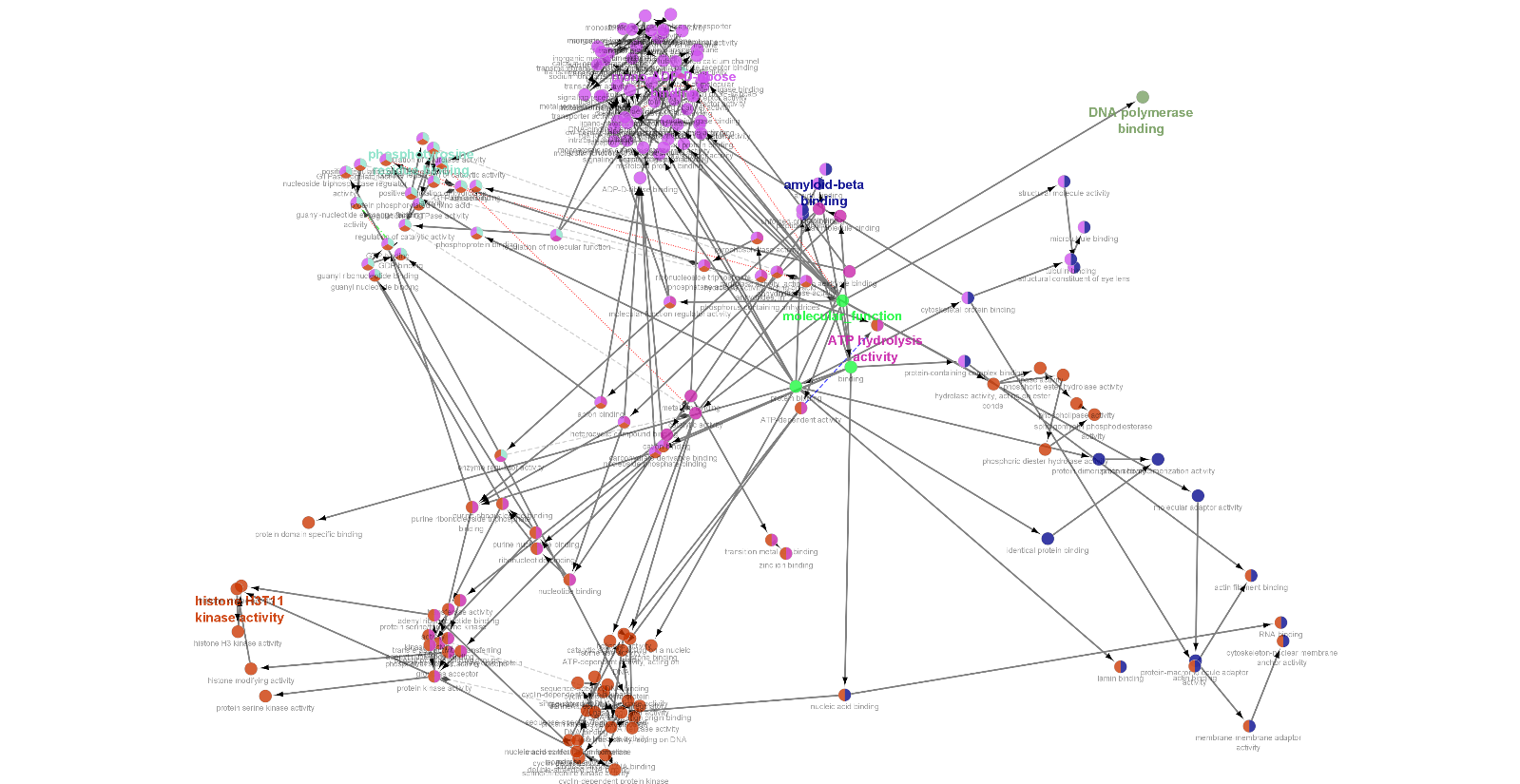

Fig. S5: Molecular functions performed by DEGs majorly highlighting DNA polymerase binding and amyloid beta binding

Pathways-

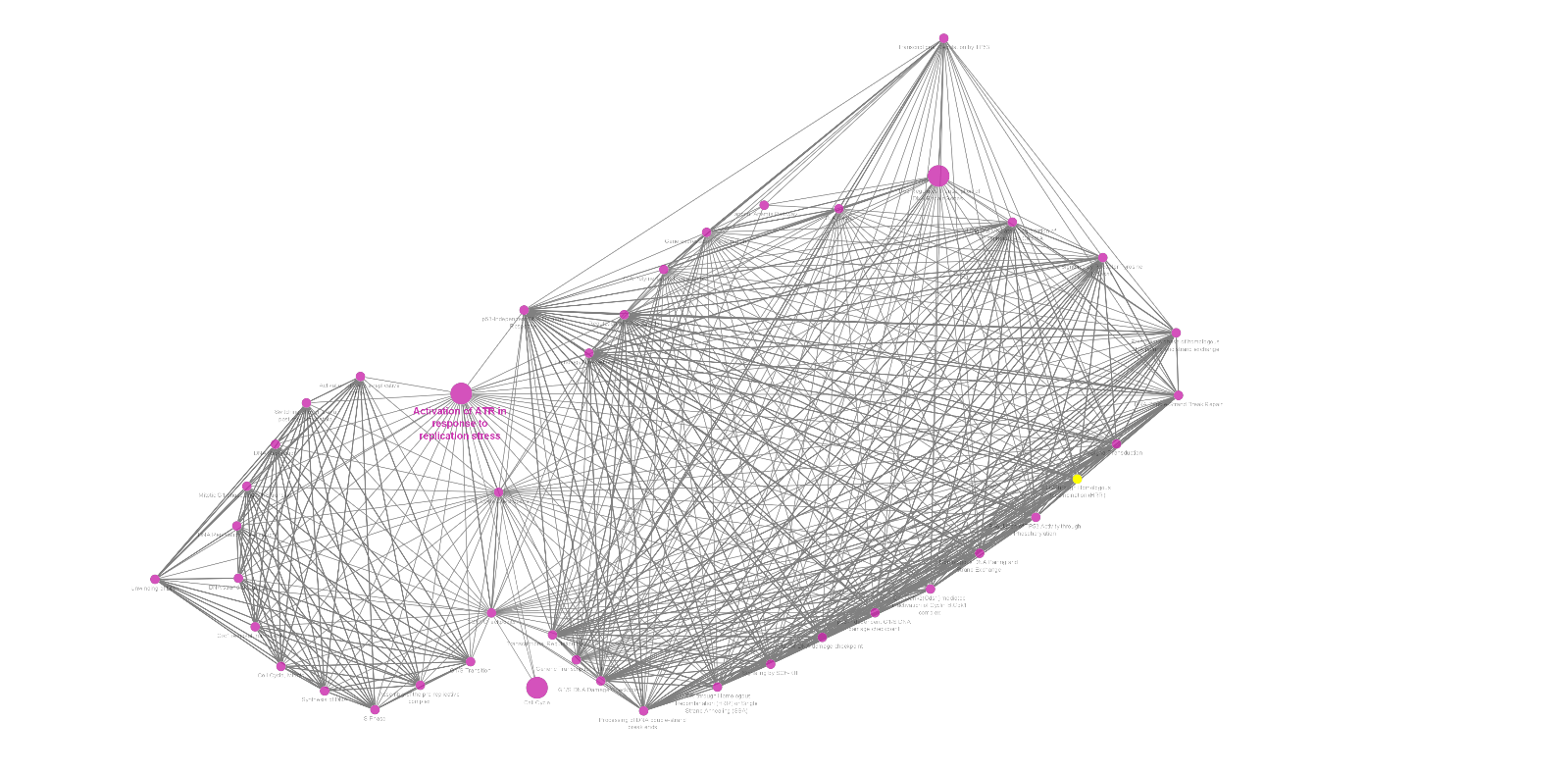

Fig. S6: Pathways associated with DEGs highlighting ATR response activation for DNA repair as the major pathway

**Stage 1 downregulated & Stage 4 upregulated genes**-

These genes were found to be aiding the tumor regression process.

Biological processes:

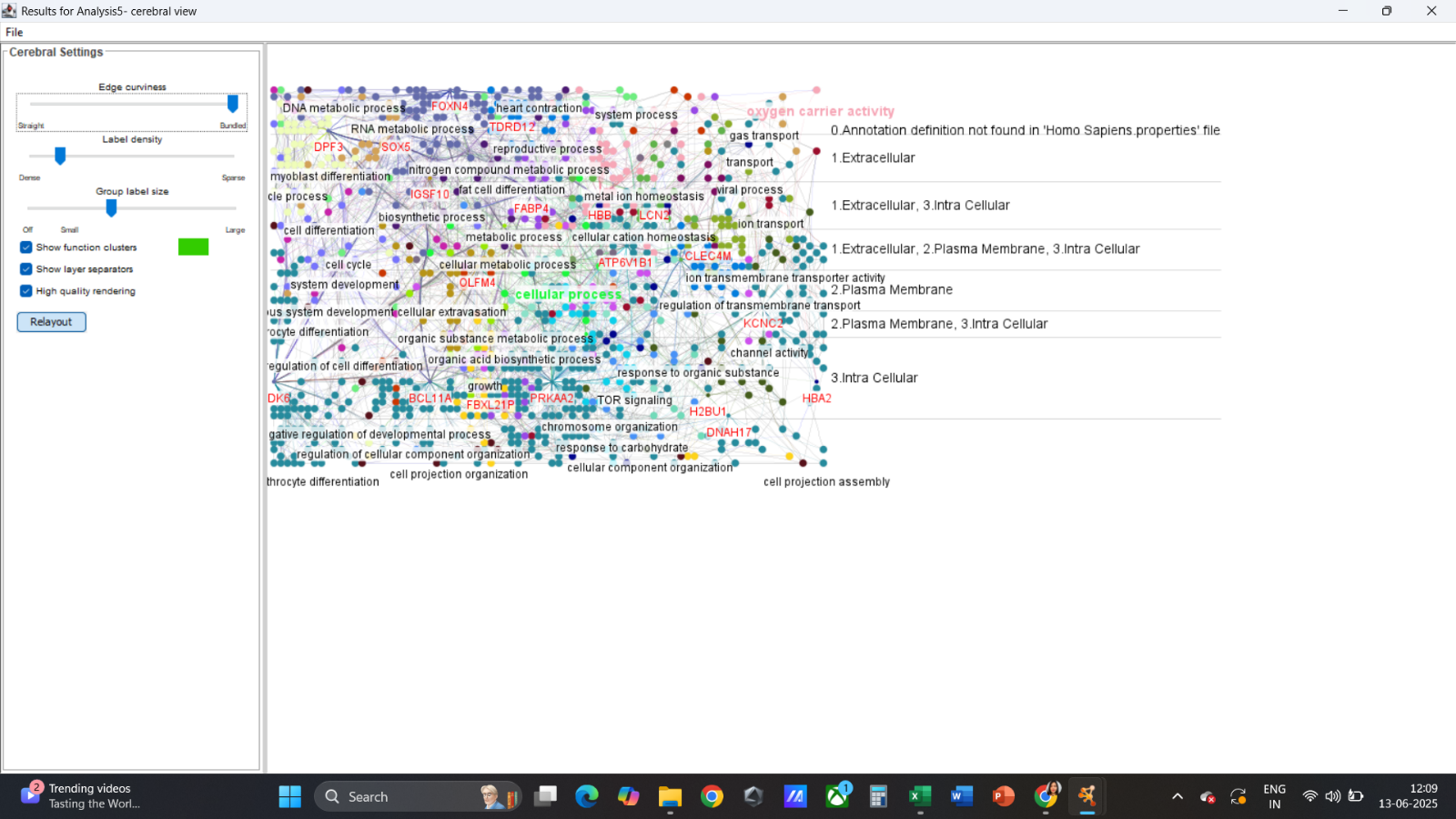

Fig. S7: shows pie chart highlighting Biological processes like microtubule organization, DNA endonuclease activity and cell-junction organization

Cellular components:

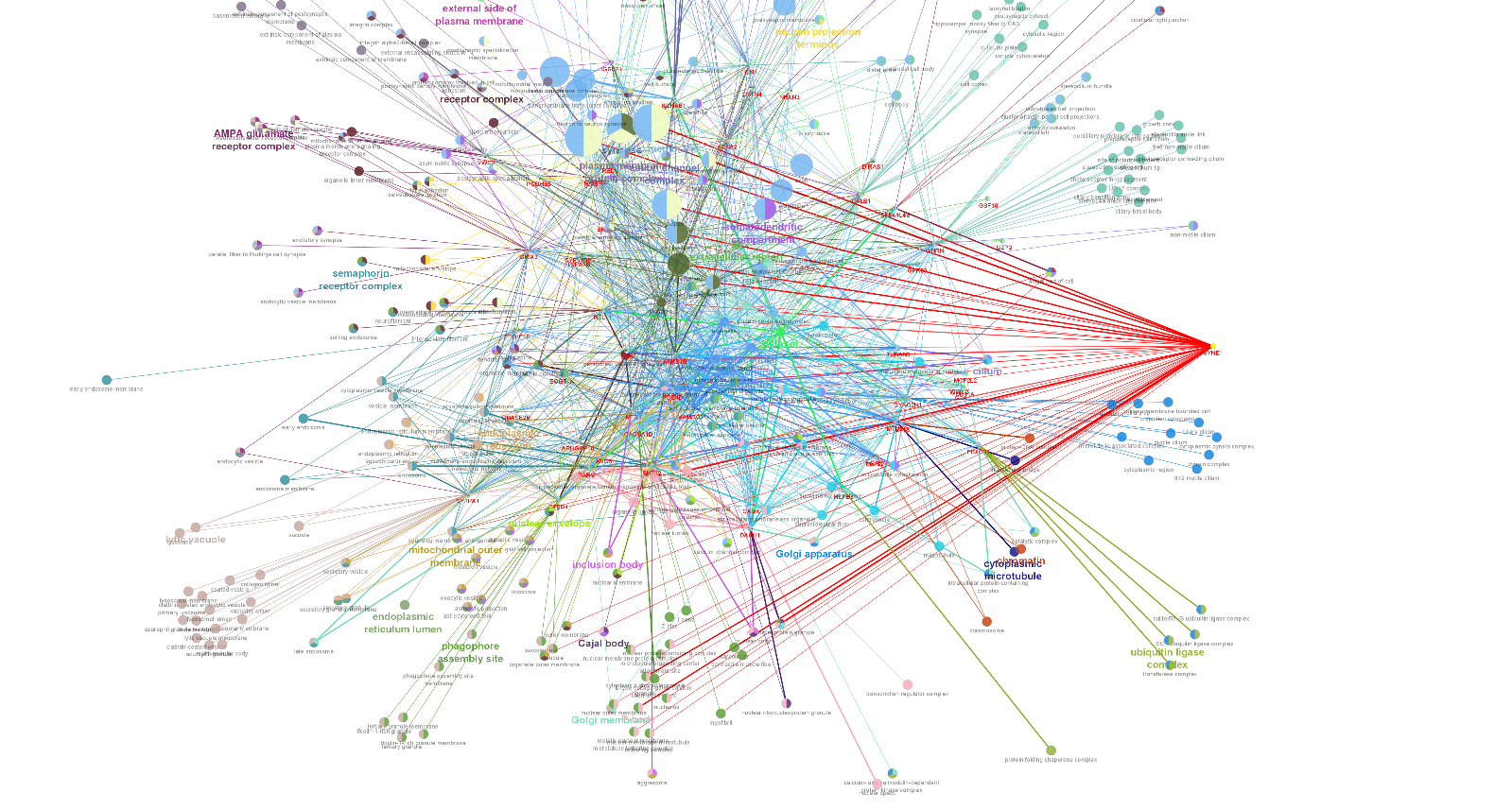

Fig. S8: shows pie chart that highlights cellular components like chromatin, nucleus and nuclear envelope

Molecular functions:

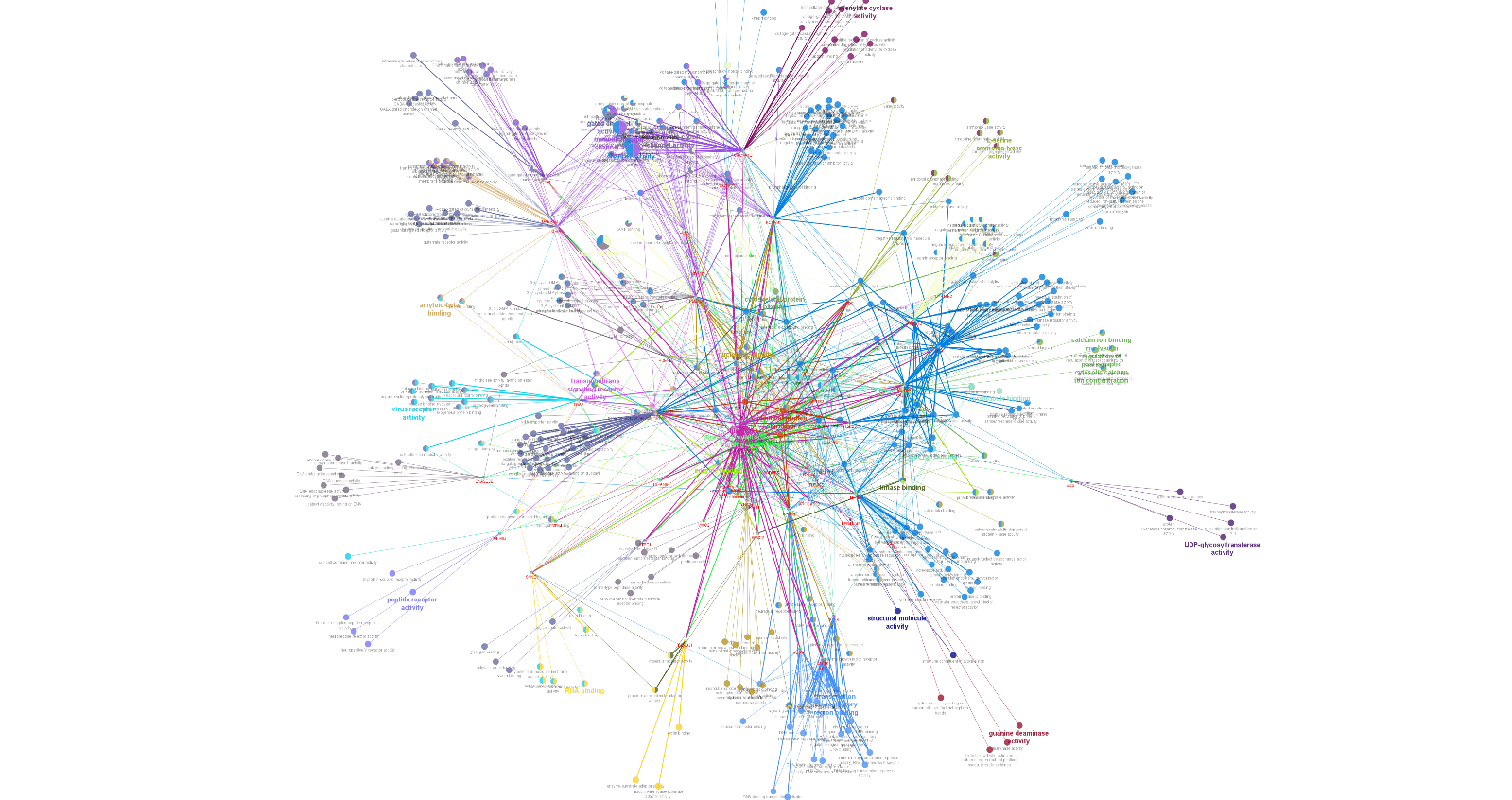

Fig. S9: shows pie chart and regulatory network highlighting molecular functions like microtubule binding, cytoskeleton protein binding

Reactome pathways:

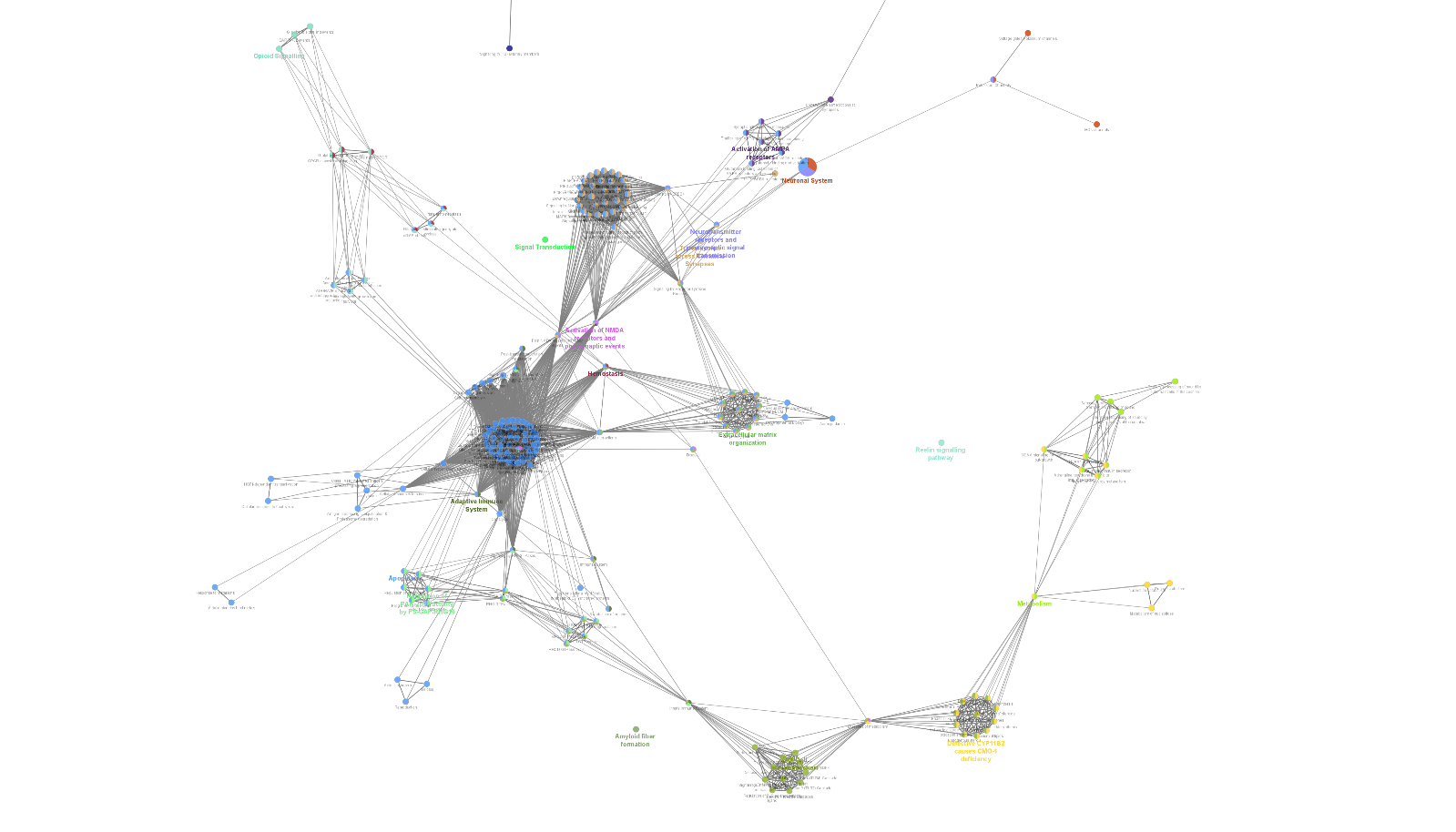

Fig. S10: shows pie chart that highlights pathways like apoptosis (for non-repairable cells) and extracellular matrix organization (repairable cells)

**Stage 1 Upregulated & Stage 4 Downregulated genes-**

These genes were found to be aiding tumor progression.

Biological processes**:**

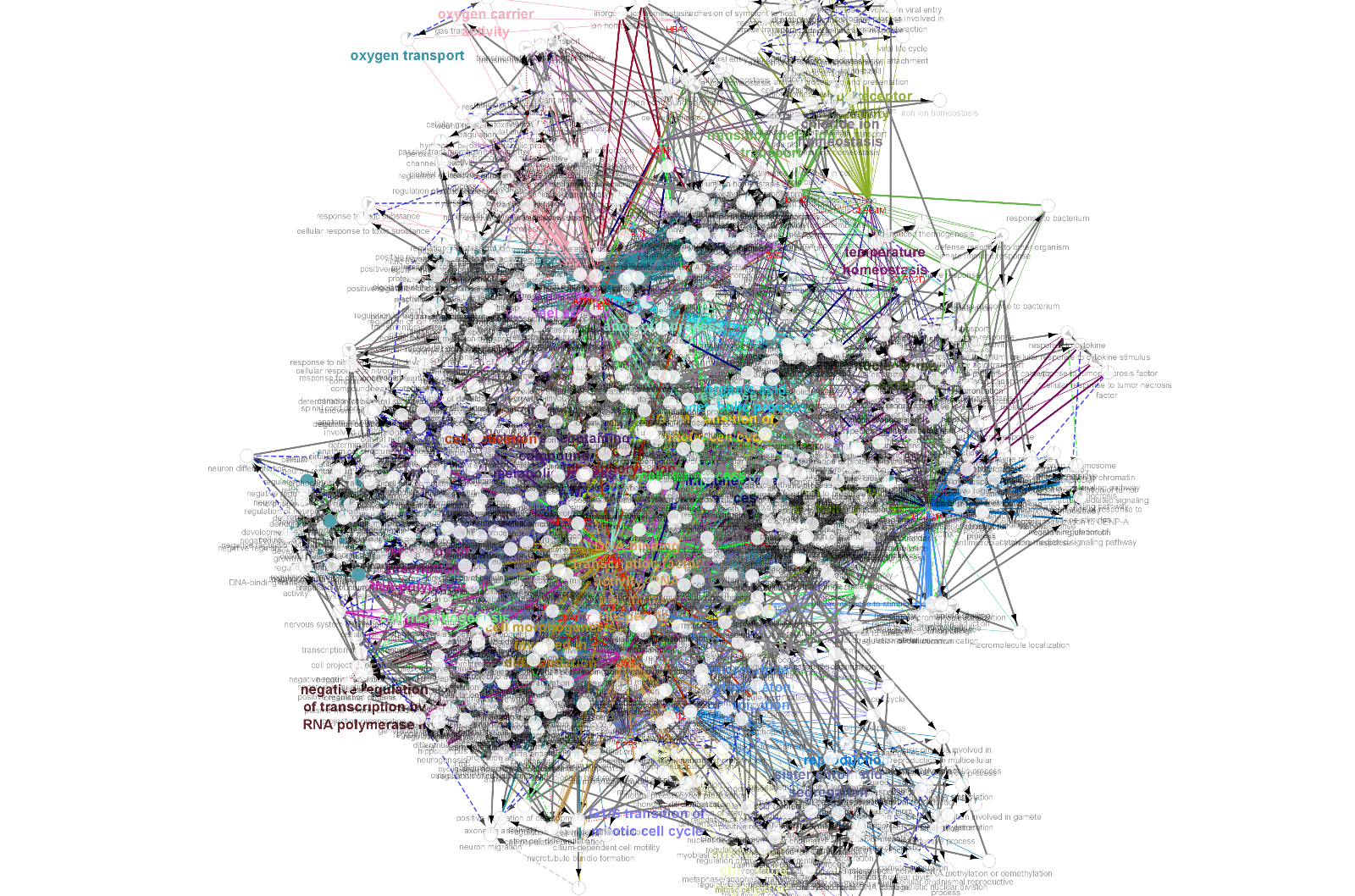

Fig. 11: shows pie chart and regulatory networks that highlight Biological processes like oxygen transport (important factor for tumor progression) and other cell division processes
